# Warming responses of microbial respiration emerge from spatially structured species interactions

**DOI:** 10.64898/2026.10.09.757985

**Authors:** Jian Wang, Ihab Hashem, Kaiyu Zhang, Satyajeet Bhonsale, Jan F.M. Van Impe

## Abstract

Microbial communities are the main drivers of global carbon cycling. Yet, predicting their respiratory responses to warming is challenging because the underlying mechanisms remain elusive. Here, we developed a spatially explicit model, integrating single-cell metabolism with metabolic exchange and endogenous heat production, to disentangle the mechanisms driving community-level temperature responses. We reveal that mutualism amplifies respiratory temperature sensitivity compared to isolated taxa, as exchanged metabolites sustain a secondary growth phase with higher respiratory demand, and this phase creates disproportionately large CO_2_ emissions under warming (apparent activation energy *E*_*a*_ increases from 0.65 to 0.77 eV). Spatial structure amplifies this further through threshold effects, whereby diffusion limitation shifts from constraining to enabling mutualistic benefits as metabolic rates increase with temperature (*E*_*a*_ increases to 1.18 ± 0.09 eV). Endogenous heat production drives additional amplification through positive feedback between local temperature rises and resource turnover rates (*E*_*a*_ further increasing to 1.78 ± 0.14 eV). Collectively, our results demonstrate that community-level temperature responses are not determined by enzyme kinetics alone but emerge from the interplay of ecological, physical, and metabolic interactions. Our model provides a general framework for understanding the combined effects of the underlying mechanisms on microbial temperature responses. Future climate–carbon models should incorporate these considerations to improve predictions.

**Graphical abstract:** 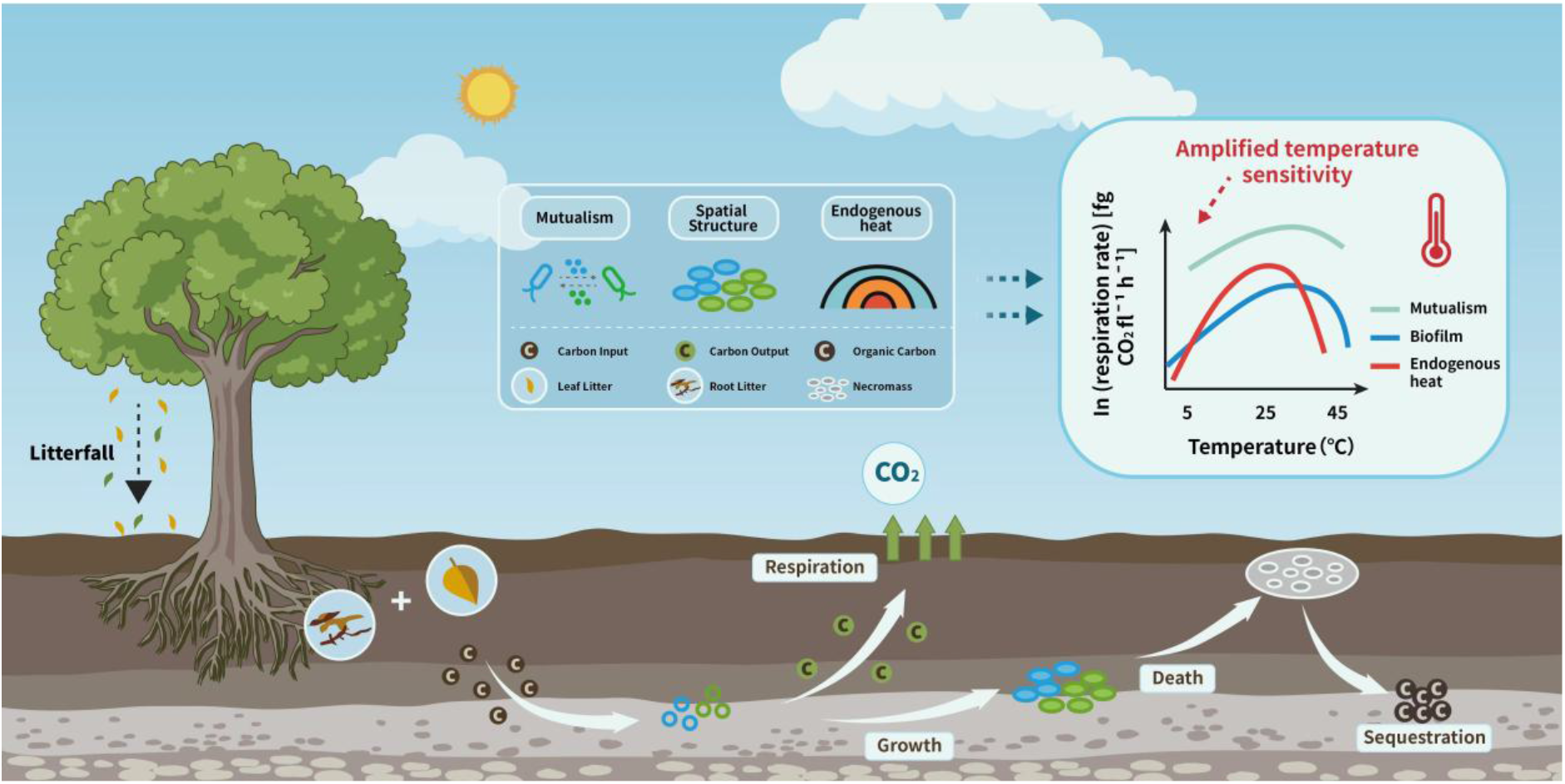

## Main

Global warming drives substantial losses of soil organic carbon (SOC)^1^, most of which is respired by microbes (1–3). Such losses further warm the atmosphere, stimulating CO_2_ emissions and reinforcing positive climate-carbon feedback (4). Therefore, understanding the mechanisms governing the microbial respiratory response to warming is critical for projecting future SOC stocks (5–7). Current approaches to this problem mainly rely on the metabolic theory of ecology (MTE) (8), which assumes that averaging temperature responses across individuals and populations adequately captures community-level dynamics (9). This assumption is effective for quantifying species-specific temperature responses, but leaves the underlying mechanisms unexplored and hinders predictions (10– 12). In particular, the role of species interactions in shaping community-level temperature responses remains poorly understood (13, 14). Species interactions occur through competitive or cooperative behaviors (15, 16). Competition is pervasive, as rapidly proliferating microbes compete for limited resources (17, 18). Microbes also engage in a variety of cooperative interactions, in which benefits such as metabolites (19, 20), vitamins (21, 22), siderophores (23, 24), or detoxification services (25, 26) provided by one species (or strain) are utilized by another, and sometimes reciprocated^2^ (27, 28). Recently, awareness has grown that warming alters these interactions. Through long-term grassland soil experiments, Yuan et al. (2021) showed that warming increases the complexity and stability of microbial interaction networks, and consequently affects the associated ecosystem processes (29). More directly, García et al. (2023) assembled bacterial communities from geothermal streams and revealed that a shift in species interactions from competition to facilitation ^3^ amplifies the temperature sensitivity of community respiration by up to 60% (30). This amplification was driven by positive feedback between species-level metabolism and biomass growth at higher temperatures. Their findings suggest that community-level warming responses are not simply the sum of individual species responses but instead emerge from temperature-dependent species interactions.

Another dimension of complexity arises from the physical structure of microbial habitats. Microbes frequently reside in spatially structured communities, such as biofilms (31). In biofilms, diffusion limitation and uneven population distributions create microhabitats that facilitate coexistence among species with different metabolic niches (32). Importantly, spatial structure sets the context for species interactions, thereby determining who interacts with whom, when, and to what degree (33). Harcombe et al. (2018) showed that when two auxotrophic bacterial species^4^ were co-cultured on solid surfaces, limited metabolite diffusion created local resource interdependence that drove the evolution of bidirectional cross-feeding (34). Similarly, Dal Co et al. (2019) showed that in two-dimensional microfluidic chambers, microscale resource gradients give rise to spatially structured glucose–acetate cross-feeding between *Escherichia coli* subpopulations. Cells near the chamber opening produced acetate, which is consumed by cells at the chamber end, where glucose was depleted (35). These results reveal that spatial structure plays an important role in regulating the strength and direction of species interactions.

In spatially structured communities, temperature directly regulates cellular metabolism and the local physicochemical environments where they interact. At the cellular level, warming shifts carbon fluxes toward higher respiratory activity and metabolite turnover. At the community level, these physiological changes modify local resource availability, thereby reshaping the strength and timing of species interactions. For example, in mutualistic consortia, warming increases both metabolite production and uptake. Partners in close proximity benefit disproportionately, as increased metabolite availability stimulates growth and respiration, creating positive feedback that amplifies the community-level temperature response. This feedback emerges within spatially structured environments, where inoculation patterns (25, 36), diffusion rates (37, 38), metabolic versatility (39, 40), and social strategies (41, 42) collectively determine the strength and timing of interactions. Yet, how these factors collectively shape community-level temperature responses remains poorly understood.

Here, we focus on a minimal two-species consortium in which species exchange secondary metabolites after consuming a shared primary substrate. This system can be viewed as a general ecological scenario in which focal populations exchange essential metabolites with neighboring species after adaptation. By integrating mutualistic interactions and endogenous heat production into a spatially explicit individual-based modeling (IbM) framework, we provide the first mechanistic account of how ecological, physical, and metabolic interactions collectively shape the temperature response of microbial community respiration. Our findings demonstrate that the interplay of these interactions gives rise to emergent community-level temperature sensitivities that substantially exceed those of isolated taxa, a result that is robust across diverse thermal performance and resource affinity scenarios. The modular framework presented here can further enable the incorporation of additional abiotic (43–46) and biotic factors (47–49), making it a valuable tool for linking microbial processes to ecosystem-scale carbon fluxes.

## Results

To explore how bottom-up mechanisms shape temperature responses of community respiration, we employed individual-based modeling (IbM) (50–52). IbM explicitly represents the traits and behaviors of individual cells, providing an integrated framework for studying the combined effects of spatial self-organization, local resource availability, and individual variability on community dynamics. It has been successfully applied to unravel ecological dynamics in bacterial colonies (53–55), biofilms (56–58), and cross-kingdom consortia (59–61). We simulated microbial growth in two cultures: (i) a well-mixed culture, in which cell positions are randomly reshuffled and nutrient concentrations are homogenized at each time period (i.e., shaken liquid culture), and (ii) a spatially structured culture, in which cells are inoculated on a hard surface and nutrients diffuse according to Fick’s law (i.e., biofilm). The model is implemented in MICRODIMS (MICRObial Dynamics Individual-based Model Simulator; (62–64). Here, we first describe the model structure and then present the results.

### Modeling the temperature response of community respiration

Our IbM considers a microbial community as a collection of expanding individual cells on a spatial grid. Cells absorb diffusing substrates from their local environments to grow, divide, produce metabolic by-products, and respire, causing the community to expand in space (Fig. 1A). Growth and respiration are coupled through bacterial growth efficiency (BGE^5^), such that a fraction of assimilated carbon is allocated to biomass growth, and the rest is respired as CO_2_ (Methods, “Carbon allocation”). BGE declines with temperature as maintenance respiration costs increase. Respiration, in turn, produces endogenous heat, which is dissipated throughout the medium. Each cell is characterized by its biomass, radius, spatial coordinates, metabolic state, and instantaneous growth rate (Fig. 1B). We postulate a minimal mutualistic consortium in which two species (species A, green; species B, blue) are exposed to a shared substrate (primary substrate, grey), and mutualistic interactions are initiated through the utilization of partner-derived metabolic by-products (secondary metabolites, orange; Fig. 1B). The shift from substrate to metabolite utilization is accompanied by a period of growth arrest, which is initiated by two conditions: (i) substrate limitation and (ii) metabolite accumulation. Once both conditions are met, the enzyme required for metabolite uptake is induced according to a temperature-dependent timer (Methods, “Nutrient uptake”; SI Appendix). Subsequently, both species rely on the partner-derived byproducts to grow (Fig. 1C). The mechanisms underlying the shift from primary substrate to secondary metabolite have been well described as ‘diauxic growth’ in laboratory cultures (66, 67). The surrounding environment is described by substrate, metabolite, and temperature fields (Fig. 1D & E).

**Fig. 1.**
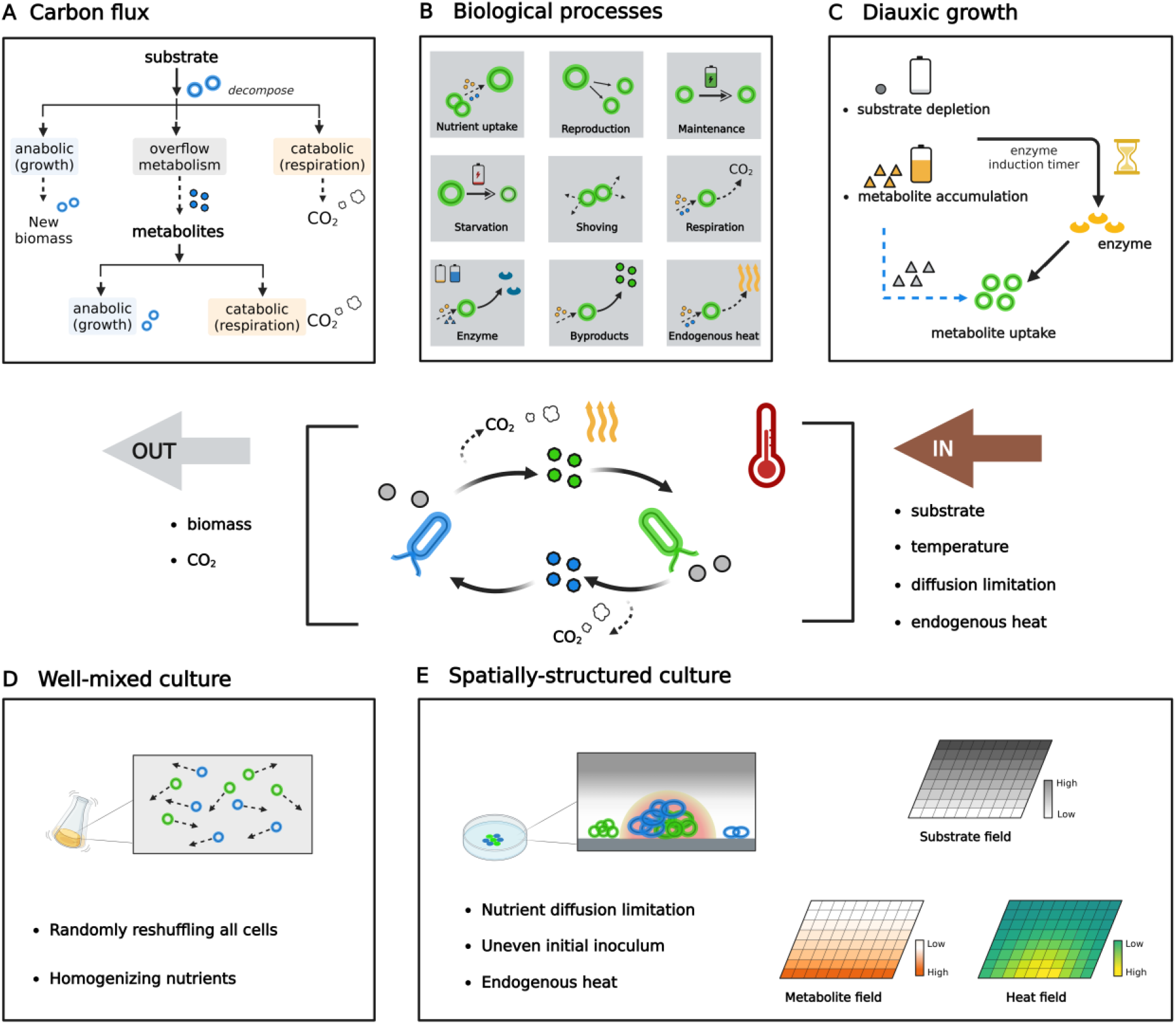
Overview of the individual-based model (IbM) for a temperature-dependent mutualistic consortium. (A) Carbon flux pathways. Two species decompose a shared primary substrate into biomass, secondary metabolites, and CO_2_. Metabolites are exchanged through cross-feeding to sustain additional biomass growth and respiration. (B) Biological processes at the cell level. Individual cells uptake nutrients to sustain maintenance, reproduction, and the production of enzymes, metabolites, and endogenous heat. If maintenance processes are not saturated by available nutrients, the deficit in nutrient uptake is compensated by degrading an equivalent amount of cellular biomass. Meanwhile, cells interact physically through shoving when spatial overlap occurs. (C) Diauxic growth mechanism. Metabolite uptake requires enzyme induction, which is triggered when two conditions are met: (i) substrate becomes limiting, and (ii) metabolite concentration exceeds a threshold. (D) Well-mixed culture. Cell positions are randomly reshuffled, and nutrient concentrations are spatially averaged at each time step to remove gradients. (E) Spatially structured culture. Cells remain fixed on the surface. Substrate, metabolites, and heat diffuse according to Fick’s law and the heat equation, respectively.

### Mutualistic interactions and carbon flux redistribution amplify the temperature sensitivity of community respiration

We first simulated the growth of microbial consortia in a well-mixed culture (Fig. 2A, left panel). To investigate the impact of mutualistic interactions on temperature-dependent respiration, we compared two scenarios: a baseline scenario, where population growth stops after depleting the primary substrate, and a mutualistic scenario, where species can utilize partner-derived metabolites after pathway switching (Fig. 2A, right panel). We found that metabolite exchange fundamentally affected community growth dynamics and subsequently altered temperature responses. When metabolite exchange is enabled, secondary metabolites sustain an additional phase of biomass production and CO_2_ emission. The left panel of Fig. 2B displays three metabolic states, which transition from substrate utilization (grey) to metabolite utilization (orange), accompanied by a period of growth arrest (red). The secondary growth phase is strongly temperature-dependent (Fig. 2B, right panel). At low temperatures, slow metabolite production and accumulation delay the onset of cross-feeding, causing the consortia to respond similarly to the baseline scenario. As temperature rises, both metabolite production and enzyme induction accelerate, allowing metabolite concentrations to reach the induction threshold earlier and sustaining a more temperature-sensitive growth phase. The right panel of Fig. 2C shows that community biomass in the mutualism scenario increases slightly with temperature and declines beyond a thermal threshold, whereas CO_2_ emissions increase sharply. This divergence arises because secondary metabolism carries higher respiratory demands, such that a greater fraction of assimilated carbon is directed toward CO_2_ production rather than biomass growth at elevated temperatures. Consequently, total community respiration increases more rapidly with temperature once metabolite exchange is active. The left panel of Fig. 2C shows that the apparent activation energy 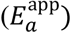 increases from 0.65 to 0.77 eV in the baseline and mutualism scenario, respectively. The developed consumer-resource model (SI Appendix, ‘*Demonstrate the outputs’*) provides the analytical basis for this finding. As warming drives the induction threshold to be reached earlier, the secondary growth phase captures a growing fraction of the total growth window, and because secondary metabolite oxidation carries a higher respiratory cost than primary substrate utilization, this increasing contribution amplifies the apparent 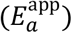 beyond the baseline enzymatic value (i.e., 0.65 eV).

**Fig. 2.**
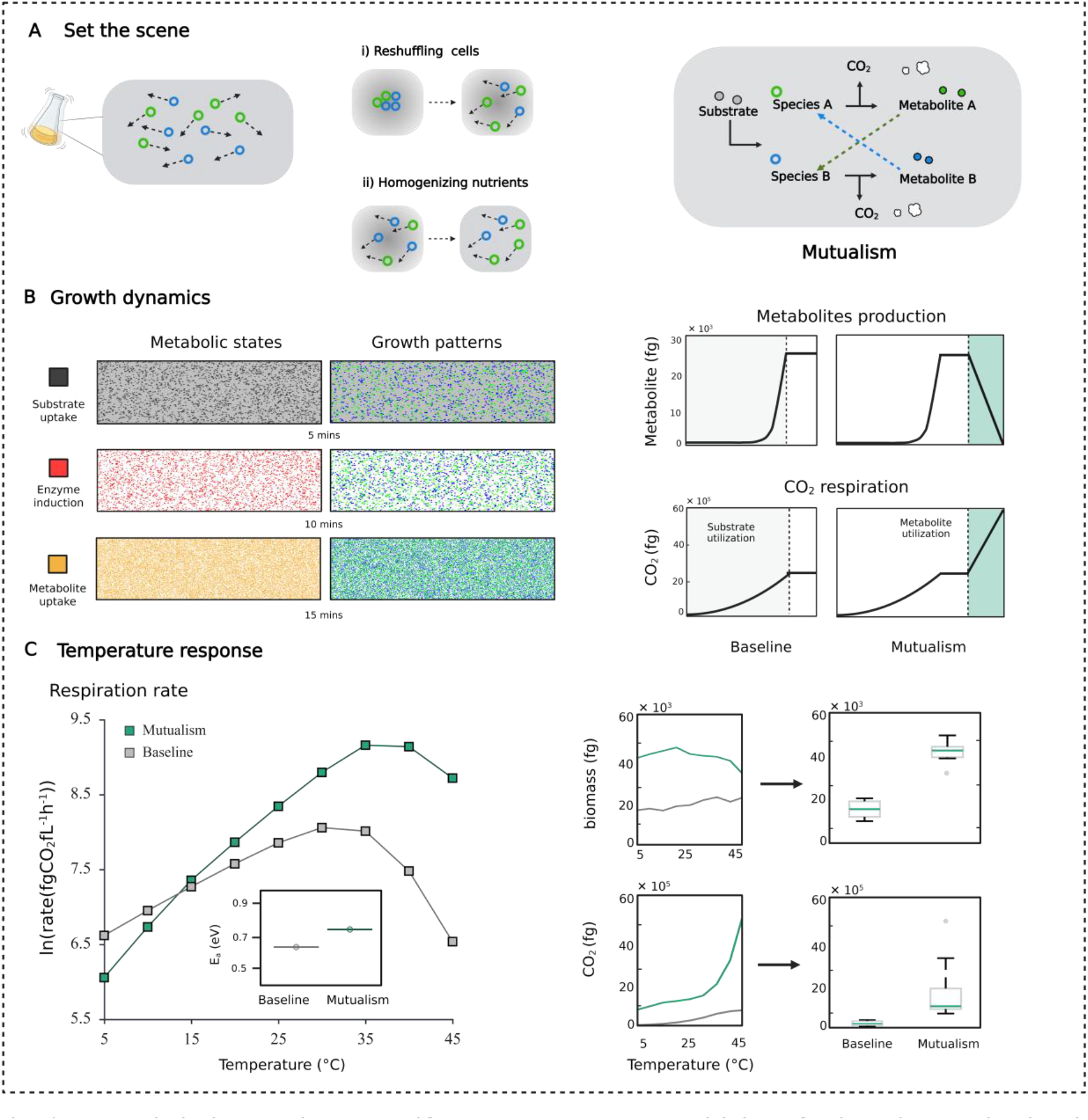
Mutualistic interactions amplify the temperature sensitivity of microbial respiration in well-mixed cultures. (A) Set the scene. Two scenarios are compared in a well-mixed culture (i.e., shaken liquid): (i) baseline, in which growth stops after substrate depletion and metabolites accumulate as waste by-products; (ii) mutualism, in which metabolite exchange is activated after substrate depletion, with each species utilizing by-products produced by its partner. In the mutualism scenario, secondary metabolites are converted into biomass and CO_2_. (B) Growth dynamics. Metabolic states of community shift from substrate utilization (grey zone) to metabolite utilization (orange zone), accompanied by a period of growth arrest (i.e., lag time, red zone). Growth patterns display the corresponding intermixing configurations of species A (green) and species B (blue). Biomass density increases over time. The lag corresponds to the enzyme induction time required to activate metabolite uptake pathways. Metabolites accumulate until enzyme induction is complete, after which metabolite utilization resumes. CO_2_ production increases sharply during metabolite consumption due to the higher respiratory demands of secondary metabolism. Example shown at 25°C. (C) Temperature response of community respiration rate. Arrhenius plots for the baseline (solid grey line, mean ± 95% CI) and mutualism (solid red line, mean ± 95% CI). Apparent activation energy 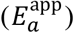 increases from 0.65 eV (baseline) to 0.77 eV (mutualism). Additional plots show that community biomass in the mutualism scenario increases slightly with temperature and declines beyond a thermal threshold, whereas CO_2_ emissions increase sharply.

### Spatial structure introduces threshold effects that amplify mutualistic temperature sensitivity

Next, we examined the effects of spatial structure on metabolite exchange. We simulated the growth of mutualistic consortia in biofilm cultures. Biofilms are highly heterogeneous, where microbes modify their surroundings through local metabolic activity. This local modification, combined with uneven population distributions, creates chemical gradients that expose cells at different positions to heterogeneous substrate and metabolite availability, thereby altering the timing and strength of mutualistic exchange.

To test this hypothesis, we repeated mutualistic simulations under biofilm conditions. Instead of reshuffling cell positions and homogenizing the nutrient field, we inoculated cells on a hard surface and allowed nutrients to diffuse (Fig. 3A, left panel). Nutrient diffusion is modeled by Fick’s second law in two dimensions. Details and nutrient field parameters are listed in the SI Appendix (Table xx). Biofilm simulations revealed two important differences compared with the well-mixed scenario. First, in well-mixed cultures, all cells experience the same substrate and metabolite concentrations and therefore follow a synchronous diauxic pattern (i.e., primary → lag → secondary). In biofilms, however, substrate depletion and metabolite accumulation occur unevenly across space. Therefore, the induction conditions for metabolic switching are met earlier in some locations. Consequently, substrate- and metabolite-based growth are simultaneously present within the same community. As shown in the left panel of Fig. 3B, at a high inoculum density, the upper layer continues to grow by utilizing substrate (grey), whereas cells in the bottom layer (green) switch to metabolite utilization when substrate becomes limiting, and metabolites accumulate. This asynchronous diauxic behavior is qualitatively consistent with the spatial self-organization of microbial metabolism observed experimentally, where local metabolic activity generates chemical gradients that partition communities into distinct zones of carbon source utilization (35, 68, 69). Second, slow diffusion makes mutualistic benefits highly dependent on partner proximity. Species A-B pairs inoculated in close proximity initiate cross-feeding earlier and accumulate more biomass, whereas spatially segregated colonies can only access partner-derived metabolites after diffusion across longer distances (snapshots are provided in the SI Appendix). This proximity dependence is qualitatively consistent with experimental observations showing that cross-feeding efficiency in microbial consortia scales with cell-to-cell distance, as short-range metabolite gradients confine mutualistic benefits to closely positioned partners (70, 71).

**Fig. 3.**
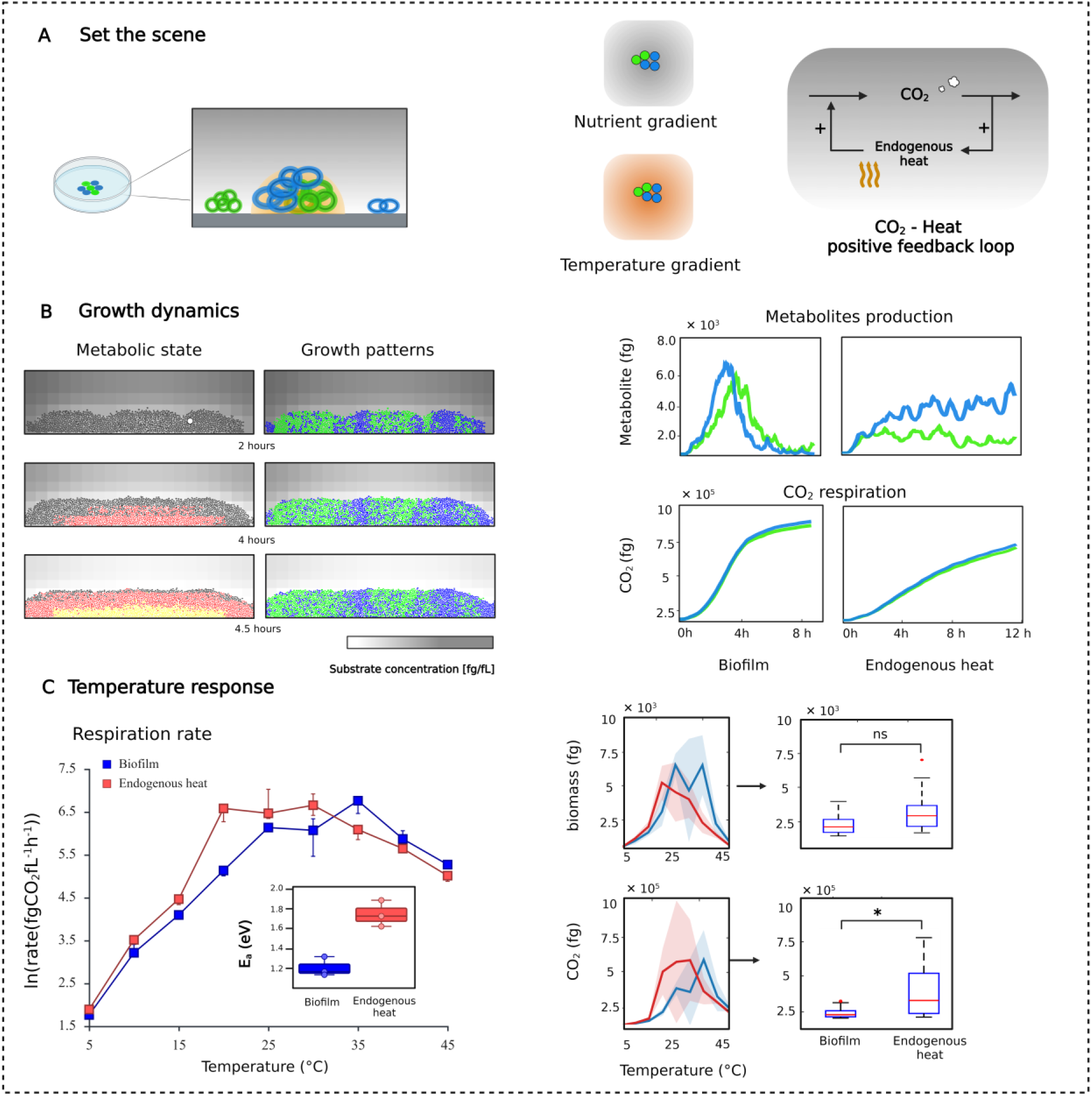
Spatial structure and endogenous heat production further amplify respiratory temperature sensitivity. (A) Set the scene. To simulate spatially structured biofilms, initial cells are fixed on a surface; substrate, metabolite, and temperature fields evolve through diffusion. Two scenarios are compared: (i) biofilm, in which nutrient gradients emerge from diffusion limitation, but temperature remains spatially uniform; and (ii) biofilm with endogenous heat, in which cells produce metabolic heat proportional to their instantaneous respiration rate, creating local temperature hotspots that feed back on metabolism. (B) Growth dynamics. Left panels: spatial biomass distributions at representation times. The community displays spatially asynchronous metabolism. The top layer continues to grow on the primary substrate (grey). The bottom layer has exhausted the local substrate and transitions to metabolite utilization (orange). The middle layer is undergoing enzyme induction (red). Growth patterns display the corresponding intermixing configurations of species A and B. Right panels: time series of community-level CO_2_ production rate (top trace) and metabolite concentration (bottom trace). Metabolite dynamics show spatial asynchrony, where metabolites are produced in substrate-rich zones and consumed in substrate-depleted zones. In the endogenous heat scenario, both CO_2_ production and metabolite concentration traces display enhanced fluctuations (right panel 2), arising from localized metabolic acceleration in temperature hotspots, where elevated local temperature simultaneously increases production and consumption rates. Statistical annotations: ns, not significant (p ≥ 0.05); *, p < 0.05 (two-sided Wilcoxon rank-sum test; n = 20 independent simulation replicates per condition). (C) Temperature response of community respiration rate. Arrhenius plots for biofilm with uniform temperature (solid blue line; *Ea* = 1.18 eV ± 0.09 eV ; 53% amplification relative to well-mixed mutualism) and biofilm with endogenous heat production (solid red line; 1.78 eV ± 0.14 eV; 131% total amplification relative to well-mixed mutualism). Shaded bands denote ±1 s.d. across simulation replicates.

Warming intensifies both of these spatial effects by accelerating resource turnover. Higher temperatures increase metabolite production and uptake rates, conferring a disproportionate growth advantage on closely packed partners relative to more distant colonies. This spatially uneven onset of mutualism produces locally favorable regions of highly active cross-feeding that dominate CO_2_ emission at elevated temperature, thereby steepening the overall temperature–respiration relationship. The left panel of Fig. 3C shows that the apparent activation energy of community respiration in biofilms exceeds that of well-mixed mutualistic consortia by approximately 53% with 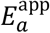 increasing from 0.77 to 1.18 ± 0.09 eV. Importantly, our simulations confirm that spatial organization does not simply slow bulk metabolic activity (72, 73). Rather, it makes mutualistic benefits increasingly temperature-dependent, as locally induced cross-feeding, reinforced by partner proximity, drives disproportionate increases in respiration at increased temperatures.

### Endogenous heat production further amplifies temperature sensitivity through local thermal feedback

Having shown that spatial structure makes mutualistic benefits increasingly temperature-dependent, we sought to further test our hypothesis by introducing endogenous heat production during cellular metabolism. Rather than imposing a spatially uniform temperature field, we coupled each cell’s metabolic activity to the local temperature (Fig. 3A, right panel). Each cell releases heat proportional to its instantaneous respiration rate, and this heat diffuses through the biofilm according to the heat equation. Thus, the local temperature changes dynamically through endogenous heat production and transfer across the spatial grid. This metabolic-temperature coupling further amplified the community response. As the biofilm grew, metabolically active clusters accumulated heat faster than isolated cells, creating local temperature hotspots where growth rates increased nonlinearly. The right panel of Fig. 3C shows that endogenous heat production (red solid line) accelerates CO_2_ emission compared to the biofilm without endogenous heat (blue solid line), allowing the pathway-switching threshold to be reached earlier within dense clusters. Accelerated pathway switching, in turn, increased local respiration and heat production, reinforcing the hotspot. At the community scale, this cascade (heat accumulation **→** accelerated cross-feeding **→** elevated CO_2_ **→** more heat) disproportionately increases total CO_2_ emission at elevated temperatures. The left panel of Fig. 3C shows that the apparent activation energy increased by a further 51%, from 1.18 ± 0.09 (biofilm, uniform temperature) to 1.78 ± 0.14 eV (biofilm with endogenous heat). Together, these three mechanisms are decomposed analytically into additive components of *E*_*a*_ amplification, enabling the relative importance of each process to be assessed (SI Appendix, Eq. 68).

### Amplification mechanisms are robust across diverse thermal and resource affinity scenarios

Our previous simulations assumed that the two species were symmetric and shared identical thermal performance curves and growth kinetics. Here, we removed this assumption to test whether the amplification mechanisms described above are robust to species-level variation. We simulated 16 scenarios by combining four thermal performance curve (TPC) hypotheses with four resource affinity scenarios (4 × 4). As in previous studies, we considered four TPC hypotheses for the two species (74): (i) hotter is better, in which the species with the higher thermal optimum also occupies a wider thermal niche; (ii) hotter is not better, in which low-temperature specialists compensate for biochemical constraints through increased enzyme efficiency; (iii) thermal trade-off, in which higher thermal optima confer greater maximum performance at the cost of reduced niche breadth; and (iv) specialist-generalist, in which one species has a narrow, high-performance thermal range and the other performs moderately across a broader range. Resource affinity scenarios were defined by varying the Monod half-saturation constants for substrate (*K*_*S*_) and metabolite (*K*_*M*_) uptake, generating four combinations of higher or lower affinity for each resource independently (Fig. 4B).

**Fig. 4.**
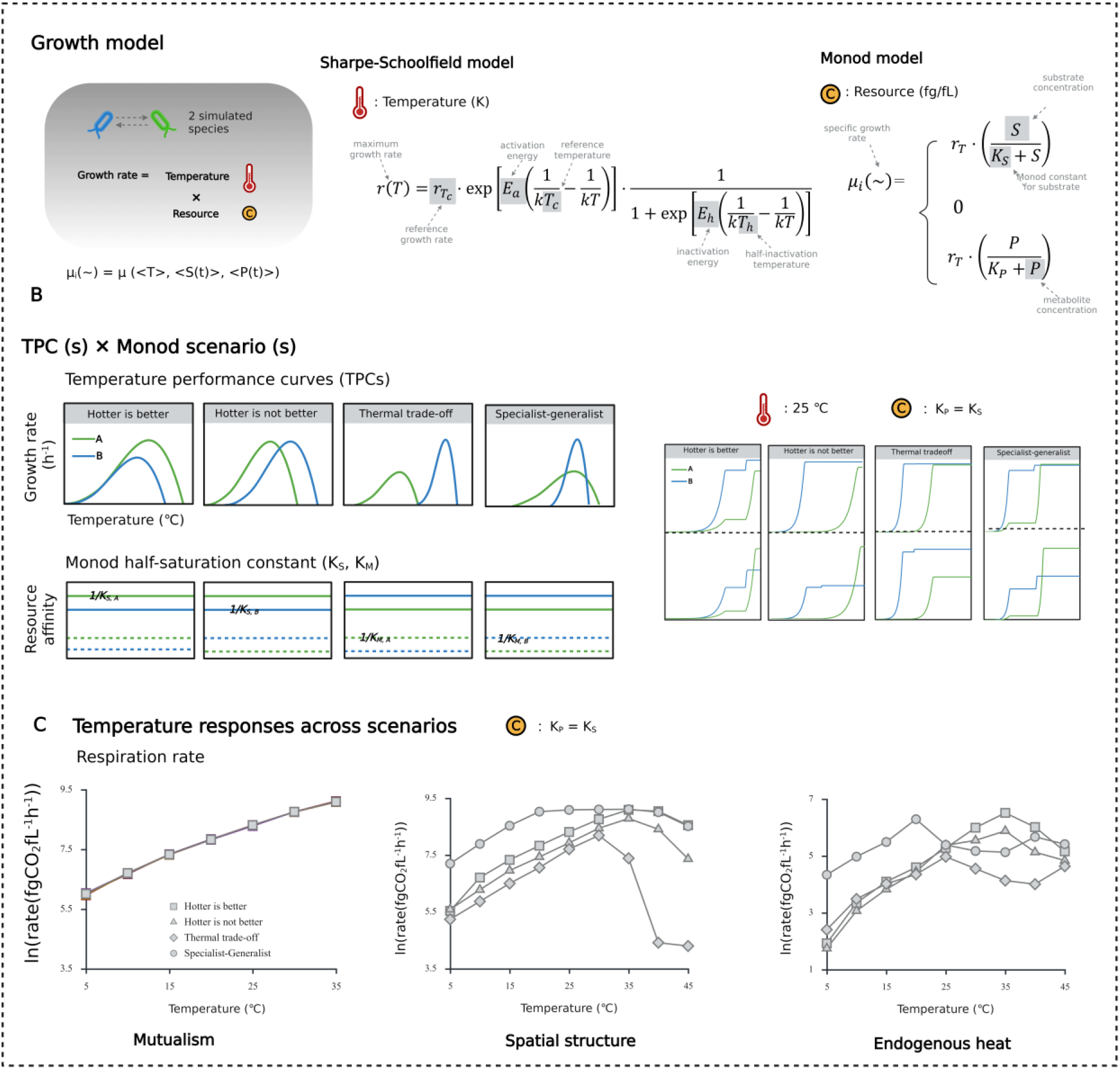
Robustness of amplification mechanisms across ecological scenarios. (A) Growth models. Individual cell growth rate is determined by temperature and resource availability, described by the Sharpe–Schoolfield and Monod models, respectively. (B) Thermal performance curve (TPC) and resource affinity scenarios. Four TPC hypotheses were tested (74): (i) hotter is better; (ii) hotter is not better; (iii) thermal trade-off; and (iv) specialist–generalist. Four resource affinity scenarios were defined by varying Monod half-saturation constants of species A relative to species B: (i) higher substrate, higher metabolite affinity; (ii) higher substrate, lower metabolite affinity; (iii) lower substrate, higher metabolite affinity; and (iv) lower substrate, lower metabolite affinity. Differences in growth kinetics create asymmetries in biomass accumulation, metabolite production, and CO_2_ respiration between species. However, the inferior competitor during primary substrate uptake can exploit partner-derived metabolites through cross-feeding, enabling stable coexistence within the mutualistic consortia. (C) Temperature response. The apparent activation energy E_a_ of community respiration was evaluated across all 16 TPC × resource affinity scenarios under the three environmental settings. No significant differences in E_a_ were observed across scenarios, confirming the robustness of the amplification mechanisms to species-level variation in thermal physiology and resource affinity.

Across all 16 combinations, differences in TPC and resource affinity introduced asymmetries during the primary substrate uptake phase. The species with faster nutrient uptake gains a biomass and metabolite production advantage (Fig. 4C). Subsequently, excess metabolic by-products from the faster-growing species are exploited by slower-growing species through cross-feeding, enabling the inferior primary competitor to contribute disproportionately to CO_2_ emission at higher temperatures. This complementary resource use stabilizes coexistence within cross-feeding consortia and preserves the amplification of temperature sensitivity across all scenarios. No statistically significant differences in apparent activation energy in exponential phase were detected among scenarios (Fig. 4C; other combinations are presented in SI Appendix ‘demonstrate the output’), confirming that the amplification mechanisms identified here are general properties of mutualistic microbial consortia.

## Discussion

We present a mechanistic viewpoint that the interplay of ecological, physical, and metabolic processes gives rise to emergent amplification of community-level temperature sensitivity in microbial respiration. Our findings are consistent with, and mechanistically explain, experimental observations of amplified CO_2_ emissions under warming in both laboratory (30) and field settings (75). Our analysis identifies three mechanisms through which this amplification occurs. The first is mutualistic cross-feeding, driven by the reciprocal exchange of essential metabolites that sustain each partner’s growth. Ample experimental evidence confirms that mutualistic interactions are common in microbial communities and expand the niche space of interacting partners (76, 77). In our model, such niche expansion occurs through the utilization of secondary metabolites after primary substrate depletion, thereby improving overall resource-use efficiency and amplifying CO_2_ emissions. Our analysis demonstrates that the onset and magnitude of secondary metabolite exchange are strongly temperature-dependent. At elevated temperatures, both metabolite production and uptake accelerate, allowing induction thresholds to be reached earlier and sustaining an efficient secondary growth phase. This shift in interaction structure from competition to mutualism under resource limitation amplifies community temperature sensitivity by approximately 20%, with *E*_*a*_ increasing from 0.65 eV to 0.77 eV.

The second mechanism is spatial structure, which regulates the strength and timing of metabolic exchange. Unlike well-mixed systems, spatially structured environments limit diffusion and create local concentration gradients of substrates and metabolites. These diffusion limitations introduce threshold effects. At lower temperatures, limited metabolite transport constrains mutualistic benefits, whereas as temperatures rise and metabolic rates accelerate, faster cluster expansion reduces the physical distance between interacting partners and enhances cross-feeding efficiency. Our analysis shows that mutualism within a spatially structured biofilm further amplifies *E*_*a*_ to 1.18 ± 0.09 eV. This finding demonstrates that the physical structure of microbial habitats, such as biofilm extracellular polymeric substances (EPS), soil aggregate structure, and pore size distribution, critically determines the temperature response of microbial communities. (73)

The third mechanism is endogenous heat production, which accelerates local resource turnover through a positive feedback between heat accumulation and enhanced metabolic activities. During nutrient oxidation, chemical energy is dissipated as heat (78). In dense, energy-rich environments where heat dissipation is relatively slow, this endogenous heat can elevate local temperatures toward or beyond the growth optimum, creating spatially heterogeneous temperature fields. Our simulations reveal that this heat-metabolism coupling fosters a self-reinforcing feedback (i.e., local warming **→** accelerated mutualistic exchange **→** elevated CO_2_ emission **→** more heat), with *E*_*a*_ increasing from 1.18 ± 0.09 to 1.78 ± 0.14 eV. Altogether, if there is sufficient understanding of the ecological mechanisms regulating microbial temperature response, the predictions of the microbe-mediated carbon cycle will be less uncertain under future climate warming (10, 65).

In reality, microbial respiratory activity is far more intricate than the description used in our simulations (2, 48, 79). Like all models, our framework makes simplifying assumptions. We consider only a minimal two-species system, assume generic secondary metabolite exchange without specifying the biochemical mechanisms (27, 80), and do not capture the evolutionary adaptation of thermal performance curves (81, 82). However, these simplifications allow biological complexity to be traded for mechanistic clarity. Rather than attempting to reproduce the exact dynamics of microbial respiration, our simulations demonstrate the community-level temperature responses that emerge when multiple ecological feedbacks are introduced to an existing modeling framework.

Our findings highlight the need for further empirical and theoretical studies. Spatially explicit experimental systems, such as microfluidic devices combined with metabolomics (83), can reveal how specific cross-feeding mechanisms respond to warming, whereas long-term warming experiments can test whether evolutionary adaptation reinforces or dampens the interaction-driven amplification we identify (48, 84). Extending our framework to multispecies communities with asymmetric interactions (52, 85, 86) and diverse metabolic cross-feeding strategies (27, 87, 88) will help determine how the mechanisms described here scale to natural soil environments.

Ultimately, we demonstrate that IbM can disentangle community-level temperature responses into their constituent ecological processes and outline a mechanistic framework for linking species interactions to climate–carbon predictions. Our results reveal that current Earth System Models (ESMs), which parameterize microbial temperature sensitivity as a fixed enzymatic property, systematically underestimate community-level warming responses. Integrating bottom-up ecological mechanisms into ESMs is essential for projecting future climate–carbon feedbacks (89).

## Methods

We employed an individual-based model (IbM) to study how ecological processes shape the temperature response of microbial respiration. By representing each microbe as a discrete entity with explicit traits and behaviors, IbM captures the effects of spatial structure, local resource availability, and individual variability on emergent community dynamics. The model dissects microbial dynamics into two coupled processes: (i) biological processes, including cell metabolism and interactions, and (ii) diffusion, governing the transport of extracellular substances such as substrates, metabolites, and heat. These processes are integrated within a hybrid Eulerian-Lagrangian framework, which couples a continuous representation of environmental gradients (Eulerian) with discrete modeling of individual cells (Lagrangian). Below, we first provide a model overview and then present the submodels for the biological and diffusion processes.

### Model overview

We considered a minimal mutualistic consortium of two species growing on a spatial grid (i.e., 40 × 20 μ*m*^2^ ). Species *A* and *B* consume the primary substrate to sustain growth, cell division, and secondary metabolite production, while simultaneously respiring CO_2_ and producing endogenous heat. The metabolic by-products *P*_*A*_ and *P*_*B*_ can only be utilized by species *B* and *A*, respectively, following a lag phase during which the required catabolic enzymes are induced. Enzyme induction is triggered by two conditions: (i) substrate limitation and (ii) metabolite accumulation. Each simulated cell is assigned a specific growth rate μ(∼) [1/h] as a function of its local microenvironment. The general expression of μ(∼) [1/h] is given by (90):

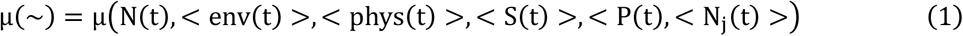

with *N*(*t*) the current cell density, < *env*(*t*) > the physicochemical properties of the environment (i.e., temperature), < *phys*(*t*) > the physiological state of cells, < *S*(*t*) > the substrate concentration,<*P*(*t*) > the metabolite concentration, and < *N*_*j*_(*t*) > the interactions with other species. These factors collectively define the microenvironment and determine cellular growth and metabolic dynamics. In turn, cells continuously modify their local environment through substrate consumption, metabolite secretion, and endogenous heat production. The interplay of these processes shapes the community’s response to environmental change.

### Biological processes (temperature dynamics, nutrient uptake, carbon allocation, metabolite yield, and interactions)

#### Temperature dynamics

The temperature dependence of the maximum growth rate follows the Sharpe-Schoolfield equation (91):

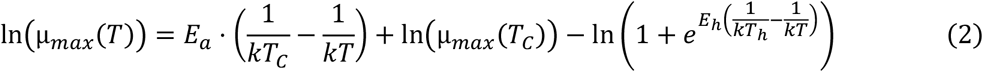

with *µ*_*max*_(*T*) [1/h] the maximum population growth rate, *k* = 8.62 × 10^−5^eV *K*^−1^ the Boltzmann’s constant, *E*_*a*_ = 0.65 *eV* the activation energy represents the temperature sensitivity of the microbial responses (i.e., the steepness of the slope of the rising part of the thermal performance curve), *T* the temperature in Kelvin (*K*), *E*_*h*_ = 2.5 *eV* the temperature-induced inactivation of growth above *T*_*h*_ = 310.15 *K* the temperature at which half of the enzymes are deactivated, and μ_max_(T_C_) = 0.8 [1/h] at reference temperature *T*_*C*_ = 293.15 *K*.

#### Nutrient uptake

Cellular nutrient uptake (substrate or metabolite) follows Monod kinetics (66):

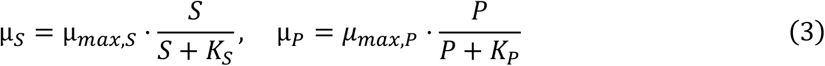

Cells initially consume substrate. The transition to metabolite utilization requires two conditions: (i) substrate limitation assessed by the supply ratio, and (ii) metabolite concentration exceeding the enzyme induction threshold (i.e., [P] > [P]^*^) (92).

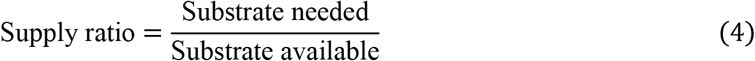

Once both conditions are met, enzyme induction begins with a temperature-dependent delay:

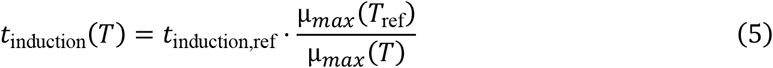

with *t*_induction, ref_ = 65 min at *T*_ref_ = 293.15 *K* .

These mechanisms for the transition of *E. coli* K-12 from glucose to acetate utilization (i.e., metabolic product) have been elucidated by Balakrishnan et al. (2021) in well-mixed cultures and Dal Co et al. (2019) in a spatially structured environment (35, 67).

#### Carbon allocation

Carbon fluxes in both metabolic pathways follow mass balance. For substrate utilization, a unit amount of substrate carbon is partitioned into biomass, metabolic byproducts, and CO_2_ according to the pathway-specific yields Y_X/S_, Y_P/S_, and 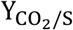, as follows:

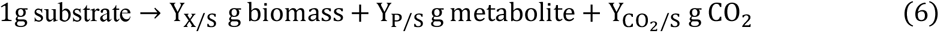

with *Y*_*X*/*S*_ = 0.45 g/g the biomass yield, *Y*_*P*/*S*_ = 0.15 g/g the metabolite yield, and 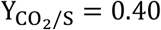 the CO_2_ yield. For metabolite utilization, carbon is partitioned into biomass and CO_2_ with yields *Y*_*X*/*P*_ and 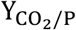, as follows:

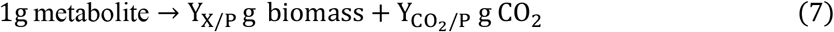

with *Y*_*X*/*P*_ = 0.29 g/g and 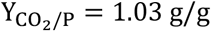, reflecting the higher respiratory cost of the metabolite. These stoichiometric yields represent the maximum anabolic conversion efficiency of each pathway. In practice, only a fraction of assimilated carbon is allocated to biomass, as a portion must satisfy maintenance respiration, which increases with temperature. To describe this constraint, we introduce a temperature-dependent bacterial growth efficiency (BGE) to account for the trade-off between anabolic growth and maintenance respiration:

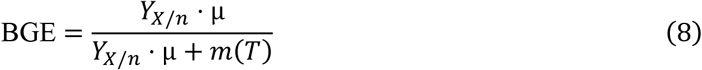

with *Y*_*X*/*n*_ the pathway-specific biomass yield, μ the potential specific growth rate, and *m*(*T*) the temperature-dependent maintenance coefficient following the Sharpe-Schoolfield model as described in Equation (2). Thus, consumed nutrients (substrate or metabolite) are first allocated to maintenance. Once maintenance requirements are met, the remaining assimilated carbon is allocated to biomass growth. This behavior is implemented using the exponential growth law that combines Monod uptake and BGE (93):

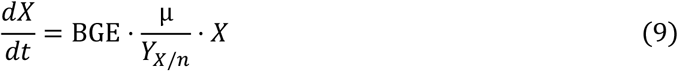

with *X* the biomass of the whole microbial population, *m*_*n*_ [fg/(fgDW.min)] the specific maintenance coefficient; *Y*_*X*/*n*_ [fgDW/fg] the cellular yield coefficient of biomass on nutrient. The remaining fraction is respired as CO_2_:

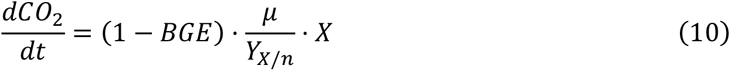

#### Respiration rate

The instantaneous community respiration rate during either phase is

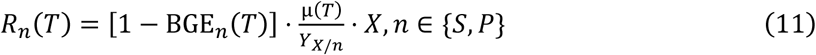

In the Boltzmann limit, 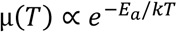 and 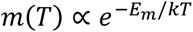. Taking the logarithmic derivative with respect to 1/*kT* yields the apparent activation energy:

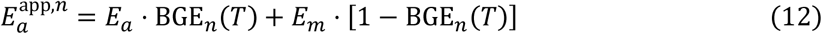

Apparent activation energy 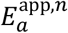 is a weighted average of *E*_*a*_ (the kinetic activation energy) and *E*_*m*_ (the maintenance activation energy), with weight [1 − BGE_*n*_(*T*)] increases with temperature as *m*(*T*) grows faster than μ(*T*).

#### Interactions

Biofilms expand by relaxing the internal pressure increase caused by biomass growth. This relaxation is effected by avoiding spatial overlap between neighboring cells, implemented using a cell-shoving mechanism (94, 95). For each cell, the vector sum of all positive overlap radii with neighboring microorganisms is calculated. The overlap radius *r*_*o*_ [μm] between a cell and its neighbor is defined as:

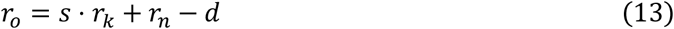

with *r*_*k*_ [μm] and *r*_*n*_ [μm] the radii of the focal cell and its neighbor, respectively, *d* [μm] the distance between their centers, and *s*[−] a scaling factor that accounts for a realistic minimum separation distance between neighboring cells.

### Diffusion processes (substrate, metabolite, and endogenous heat)

#### Substrate and metabolite diffusion

Substrate and metabolite diffusion are modeled by Fick’s second law in a 2D isotropic medium, which is expressed as follows:

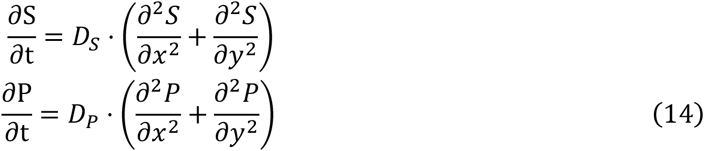

with *S* and *P* [fg/fL] the substrate and metabolites concentrations, respectively. *D*_*S*_ and *D*_*P*_ [μm^2^/min] the diffusion coefficient of substrate and metabolite, respectively. *x* [μm] and *y* [μm] the two spatial dimensions, and *t* [min] the time.

#### Metabolic heat production and transfer

Endogenous heat production per cell is assumed to be proportional to its metabolic rate:

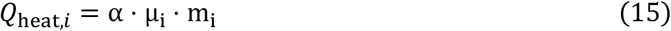

with α [K.min.*fg*^−1^] the metabolic heat coefficient, μ_*i*_ the specific growth rate and *m*_*i*_ the biomass of cell *i*, respectively. The heat transfer is modeled by the heat equation:

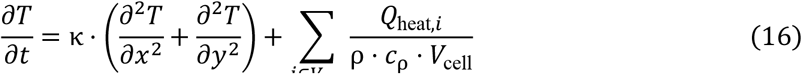

with κ [m^2^/s] the thermal diffusivity; ρ ⋅ *c*_ρ_ [J/ (m^3^ ⋅ K)] the volumetric heat capacity, and *V*_cell_ the grid cell volume.

## Data and code availability

All data and code used to replicate the figures, analyses, and findings in this paper can be found at:

## Competing interests

The authors declare no competing interests.

## Funding

This work was supported by the European Union’s Horizon 2020 research and innovation program “E-MUSE” under the Marie Skłodowska-Curie grant agreement No. 956126 (applied to J.W. and J.F.M.V.I); the European Union’s Horizon Europe research and innovation program “SUPREME” under the grant agreement No. 101058422 (applied to I.H. and J.F.M.V.I); and the European Commission within the framework of the Erasmus+ program ‘FOOD4S’ 619864-EPP-1-2020-1-BE-EPPKA1-JMD-MOB (applied to K.Z. and J.F.M.V.I).

## Author contributions

All authors conceptualized the study. J.W., K.Z., and I.H. designed the simulation. J.W. performed the study and wrote the manuscript. I.H. and J.F.M.V.I. supervised the work. J.F.M.V.I. provided funding support. All authors edited the manuscript and approved the final version. Authors’ ORCID:

## Acknowledgments

The authors thank Chen Sun (ACTA, University of Amsterdam) and Dr. Francesco Giglio (School of Mathematics and Statistics, University of Glasgow), Dr. Lydia Katsini (BioTeC+, KU Leuven) for constructive comments on an earlier version of the manuscript.

## Additional information

Supplementary material

## Footnotes

1 Soil organic carbon refers to the living and dead biological materials of soils, such as plant residues, microbial biomass, and humified organic matter.

2 Microbial beneficial interactions can be classified by two dimensions: i) the degree of reciprocity and ii) the investment of the interacting partners (27). Here, we focus on a minimal two-species mutualistic consortia.

3 We use ‘facilitation’ rather than ‘cooperation’ here to reflect the terminology of the original study (30). Authors assembled an eight-species community and characterized emergent community-level thermal responses, without identifying specific pairwise cooperative relationships.

4 Auxotrophic refers to species that cannot synthesize certain essential compounds for growth.

5 Refers to the fraction of organic carbon allocated for biomass growth to the organic carbon taken up. BGE increases as more carbon is allocated to growth than to respiration, and BGE declines when respiratory costs rise (65).

## Notes

https://gitlab.kuleuven.be/u0154531/warming-microbial-respiration

### Competing Interest Statement

The authors have declared no competing interest.

