## Supplementary material for "Warming responses of microbial respiration emerge from spatially structured species interactions"

### Warming response of microbial respiration emerges from spatially structured interactions

#### This PDF file includes:

Materials and Methods

Figs. S1 to S12

Tables S1 to S4

References

#### Material and Methods

The individual-based model (IbM) developed in this study follows the 5D protocol (*1, 2*). We first define the problem by specifying the biological question, entities, state variables, and spatiotemporal scales. We then dissect the system into the sub-processes governing microbial growth and nutrient transport, and present their population-level formulations. Next, we develop the model by discretizing each sub-process into individual cell-level. We deploy the simulation by specifying initialization procedures, boundary conditions, and process scheduling. Finally, we demonstrate the outputs by comparing IbM results against analytical predictions and comparing outcomes across ecological scenarios.

##### Define the problem

*Objective.* Our IbM aims to investigate how ecological processes shape the temperature response of microbial respiration, focusing on three mechanisms: (i) mutualistic interactions, (ii) spatial structure, and (iii) endogenous heat production. Mutualistic interaction refers to the reciprocal exchange of essential metabolites between interacting partners. Experimental evidence has demonstrated that mutualistic interactions are widespread and underpin the stability and function of microbial communities (*3, 4*). Recent experimental work further suggests that such mutualistic interactions can amplify the temperature sensitivity of community respiration (*5*). Spatial structure arises when microbes self-organize into aggregates or biofilms on surfaces (*6*). This self-imposed structure restricts nutrient diffusion, creating local concentration gradients of substrates and metabolites. Here, we focus on biofilm-type spatial organization. Endogenous heat production occurs when a fraction of chemical energy released during nutrient oxidation is dissipated as heat rather than conserved in biomass or ATP. In dense, energy-rich environments where heat dissipates slowly, this metabolic heat can raise local temperatures toward or beyond the growth optimum, creating spatially heterogeneous temperature fields that feed back on metabolic activity (*7*). Although substantial *in situ* and *in vitro* evidence highlights the ecological importance of each mechanism individually (*8–11*), how they act separately and collectively to shape community-level temperature responses remains poorly understood (*12*). Thus, we want to integrate these mechanisms to quantitatively and systematically understand their effects. Given that microbial respiration drives soil carbon turnover, a more accurate characterization of these ecological amplification mechanisms could directly inform Earth System Models (ESMs) used to project climate–carbon feedbacks under warming (*13–15*).

*Set the scene: entities, state variables, and scales.* Our IbM consists of two types of entities: individual microbial cells and a discretized environmental grid. Each cell is characterized by the following state variables: biomass $m_i$ [fg], radius $r_i$ [$\mu$m], spatial coordinates $(x_i, y_i)$ [$\mu$m], metabolic state (substrate uptake, enzyme induction, or metabolite uptake), and specific growth rate $\mu_i$ [1/h]. The environmental grid is characterized by local concentrations of primary substrate $S$ [fg/fL], secondary metabolites $P$ [fg/fL], and temperature $T$ [K]. The spatial domain is a $40 \times 20$ $\mu$m$^2$ grid with a resolution on the order of individual cell size (~2 $\mu$m). Temporal scales span from minutes (nutrient uptake, diffusion) to hours (biomass growth, cell division), resolved by a dual time-step scheme: $\Delta t_1 = 0.0005$ min for fast processes and $\Delta t_2 = 0.1$ min for slow processes (*16*). We simulated temperatures ranging from 5 to 45 °C in increments of 5 °C.

#### Dissect the system

Our IbM conceptualizes the microbial community response to temperature through the interplay of biological and diffusion processes (*17*). Biological processes describe how individual cells sense their environment, acquire resources, allocate carbon, exchange metabolites, and interact physically. Diffusion processes describe how substrates, metabolites, and endogenous heat are transported across the spatial grids. The population growth rate was determined by the combined effects of nutrient availability (i.e., $S(t)$ and $P(t)$), temperature (i.e., $T$), the physiological state of the cells (i.e., phys$(t)$), and interactions with other species (i.e., $N_j(t)$) as expressed in the following general form (*18*):

$$\mu(\sim) = \mu(N(t), < env(t) >, < phys(t) >, < S(t) >, < P(t) >, < N_j(t) >) \tag{1}$$

Our IbM adopts an object-oriented programming framework in which each population is discretized into a bunch of individual cells (*19*). Each cell is assigned a growth rate based on the local microenvironment it inhabits. In this subsection, we first dissect the microbial growth into multiple sub-processes and then present its population-level formulation.

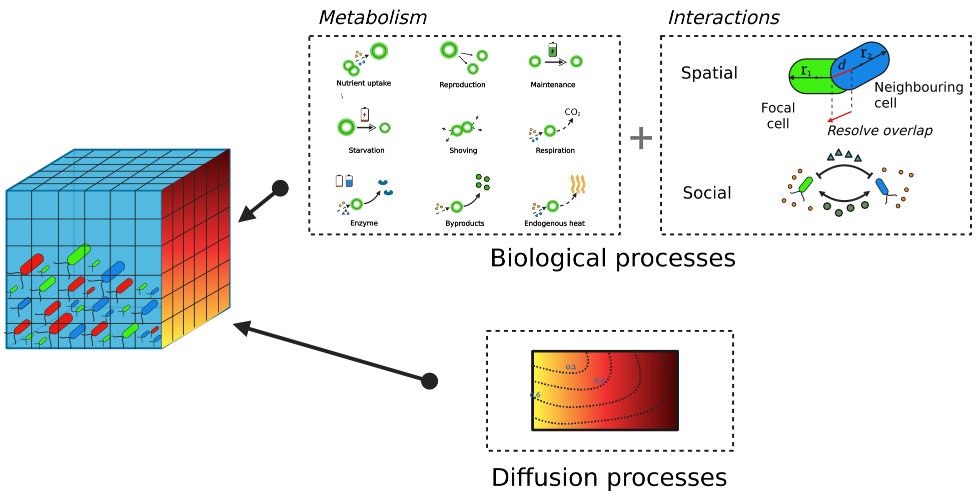


**Figure 1**: **Biological and diffusion processes in IbM**

*Temperature-dependent growth.* Temperature is a fundamental physical constraint on microbial life (*20*). As the temperature rises, the rate of chemical and enzymatic reactions in the cell increases, raising the growth rate until an optimum is reached. Beyond this optimum, further warming causes the growth rate to decline and eventually cease. This unimodal response, known as the thermal performance curve (TPC) (*21*), arises from two mechanisms that act together across the physiological temperature range. The first is thermodynamic activation. Enzymatic reactions are rate-limited by the energy barrier separating reactants from products. Temperature determines the fraction of enzyme–substrate complexes with sufficient thermal energy to cross this barrier. In the sub-optimal range, increasing temperature raises that fraction, accelerating reaction rates in a way described by Arrhenius kinetics (*22*). The steepness of this acceleration is set by the activation energy $E_a$ [eV]. A higher $E_a$ means a steeper response to warming. Across microbial taxa, $E_a$ for respiratory metabolism is approximately 0.65 eV (*23*). This value is the enzyme-kinetic baseline against which ecological amplification is measured in this study. The second is thermal inactivation. Above a species-specific optimum, rising temperature denatures the enzymes driving growth. Inactivation becomes significant once the temperature approaches $T_h$ [K], the temperature at which half of the enzymatic capacity is lost. The sharpness of this decline is described by the inactivation energy $E_h$ [eV]. A higher $E_h$ means a more abrupt drop near $T_h$. The combined effects of these two mechanisms on the maximum growth rate can be written in a separable form:

$$r(T) = r_{T_C} \cdot \underbrace{f(E_a, T, T_C)}_{\text{activation term}} \cdot \underbrace{g(E_h, T_h, T)}_{\text{inactivation term}} \tag{2}$$

Activation term $f(E_a, T, T_C)$ describes Arrhenius acceleration relative to a reference temperature $T_C$ [K] at which the growth rate $r_{T_C}$ [1/h] is defined as the reference growth rate. Mathematically, the inactivation term $g(E_h, T_h, T)$ describes the suppression as a function of how far $T$ deviates from $T_h$.

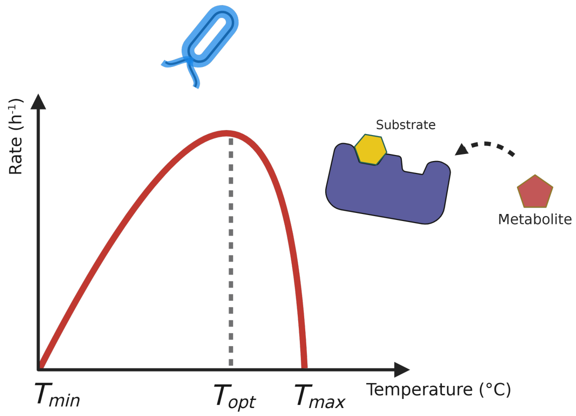


**Figure 2**: **Thermal performance curve of microbes**

*Resource-dependent growth.* The growth rate of a microbial population also depends on the availability of nutrients in the local environment. Cells take up dissolved nutrients from their surroundings and convert them into biomass. As nutrient concentration rises, the uptake rate increases, but this increase is not proportional. At low concentrations, small increases in nutrient availability produce large gains in growth rate, whereas at high concentrations, the system saturates and growth rate approaches a maximum. This hyperbolic relationship between nutrients concentration and growth rate is the defining feature of nutrient-limited microbial growth. In our system, cells can grow on either a primary substrate $S$ or a secondary metabolite $P$ produced by partner species. The nutrient-limiting relationship is described by the Monod model (*24*). The half-saturation constant $K_S$ [fg/fL] sets the substrate concentration at which the growth rate reaches half its maximum. The population with a lower $K_S$ reaches saturation at lower concentrations and is thus more competitive under substrate limitation. The same principle applies to metabolite uptake, set by $K_P$ [fg/fL]. The pathway-specific maximum growth rate $\mu_{\max}$ [1/h] is the maximum growth rate once nutrients are saturating.

$$\mu(\text{nutrient}) = \mu_{\text{max,nutrient}} \cdot \frac{\text{nutrient}}{K_{\text{nutrient}} + \text{nutrient}} \tag{3}$$

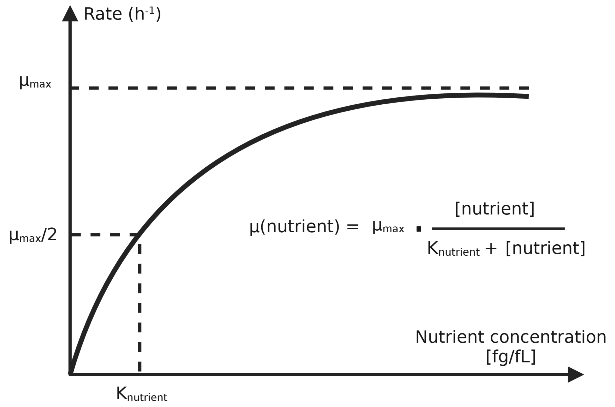


**Figure 3**: **The effect of resource availability on growth and Monod kinetics**.

*Metabolite production and respiration.* When a cell takes up a carbon source, the assimilated carbon is not entirely converted to biomass. It is partitioned among three pathways: new biomass, secreted metabolic byproducts, and $CO_2$ released through respiration (*25*). The fraction directed to each pathway is set by the biochemistry of the metabolic pathway used. During primary substrate utilization, cells perform incomplete oxidation, so that a portion of the substrate carbon is excreted as organic metabolites rather than fully oxidized to $CO_2$. This overflow metabolism arises because the biosynthetic flux exceeds the cell's capacity to fully channel carbon through central catabolism (*26*). The yield coefficient $Y_{X/S}$ [fg/fg] describes the amounts of substrate carbon fixed into biomass, $Y_{P/S}$ [fg/fg] describes the amounts excreted as metabolite, and $Y_{CO_2/S}$ [fg/fg] the amounts respired as below:

$$1 \text{ fg substrate} \rightarrow Y_{X/S} \text{ fg biomass} + Y_{P/S} \text{ fg metabolite} + Y_{CO_2/S} \text{ fg } CO_2 \quad (4)$$

During secondary metabolite utilization, no further metabolite is excreted. The partner species consumes the accumulated metabolite and oxidizes it more completely, resulting in a lower biomass yield $Y_{X/P}$ [fg/fg] and a higher respiratory cost $Y_{CO_2/P}$ [fg/fg] compared to the substrate pathway:

$$1 \text{ fg metabolite} \rightarrow Y_{X/M} \text{ fg biomass} + Y_{CO_2/M} \text{ fg } CO_2 \quad (5)$$

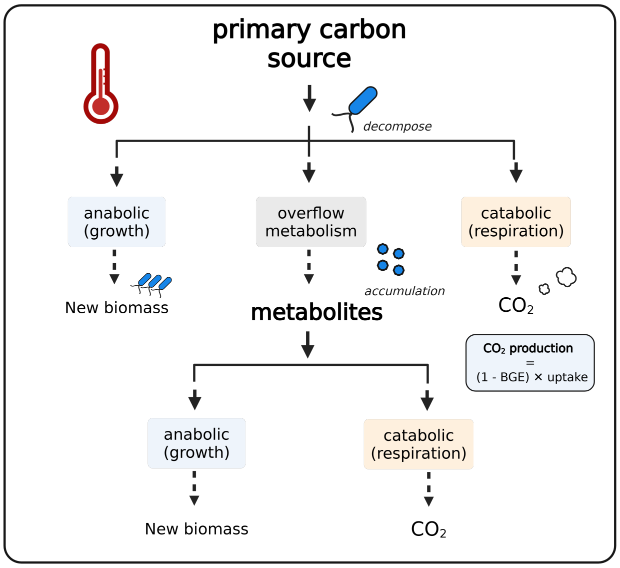


**Figure 4**: **Carbon flux**

The yield coefficients above represent the maximum anabolic conversion under resource-saturating conditions. In practice, cells must continuously expend energy to maintain cellular integrity regardless of whether they are actively growing. This maintenance demand is met by diverting a fraction of assimilated carbon away from biosynthesis and toward respiration. The bacterial growth efficiency (BGE) quantifies the fraction of assimilated carbon that is ultimately directed to new biomass (*27*). It is set by the balance between the growth rate $\mu$ and the maintenance coefficient $m(T)$. When growth is fast relative to maintenance, most carbon goes to biomass, and BGE is high; when maintenance is large relative to growth, more carbon is respired, and BGE is low. Critically, maintenance costs increase with temperature faster than the growth rate does, because the activation energy of maintenance metabolism $E_m$ exceeds that of biosynthesis $E_a$. As a result, BGE decreases with warming, and a progressively larger fraction of assimilated carbon is lost as $CO_2$ at elevated temperatures (*28*). The general form of BGE as a function of growth rate and maintenance is:

$$\mathrm{BGE}(T) = h(\mu(\sim, T), m(T)) \tag{6}$$

with $h$ the function that increases with $\mu(\sim, T)$ and decrease with $m(T)$. The community-level $CO_2$ production rate then follows directly from BGE. The fraction of carbon not retained in biomass is respired, so:

$$\frac{dX}{dt} = \mathrm{BGE}(T) \cdot v \tag{7}$$

$$\frac{d\mathrm{CO}_2}{dt} = (1 - \mathrm{BGE}(T)) \cdot v \tag{8}$$

with $v$ the carbon flux from nutrient to cell.

*Diauxic switching.* When multiple carbon sources are available, microbial cells do not consume them simultaneously. They preferentially consume the carbon source that supports the fastest growth and suppress the uptake of inferior resources until the preferred source is depleted (*24*). This sequential utilization arises from two coupled regulatory mechanisms. The first is catabolite repression. When the preferred substrate is present, intracellular signaling actively represses the transcription of genes encoding the catabolic enzymes for secondary carbon sources. The secondary pathway is not merely outcompeted in kinetics—it is transcriptionally inactivated. Only when the preferred substrate falls below a critical threshold does repression lift, and the secondary pathway becomes accessible (*29*). The active metabolic state of the population can thus be represented by a discrete pathway indicator:

$$I^S = \mathbf{1}\{\mu(\sim, S) > \mu(\sim, P)\}, \quad I^P = 1 - I^S \tag{9}$$

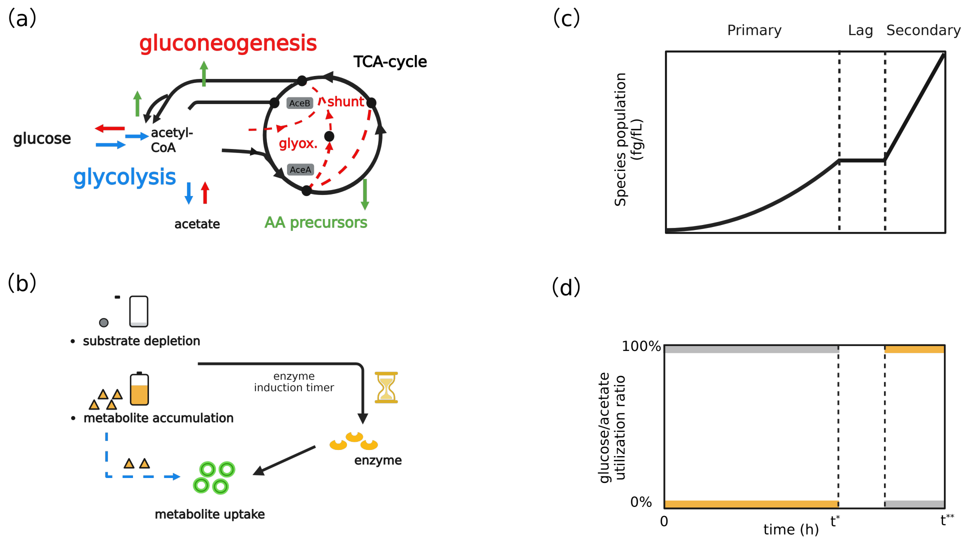


**Figure 5**: **Diauxic switching mechanisms**.

The switch is not instantaneous, however. Even after repression lifts, cells must synthesize the enzymes required for secondary carbon uptake. This enzyme induction takes time, as the cell must transcribe the relevant genes, translate the mRNA, and accumulate sufficient enzyme to sustain growth on the new substrate (*29, 30*). The induction lag $t_{\text{ind}}$ [min] represents this delay between the environmental signal (substrate depletion) and the physiological response (onset of secondary growth). Critically, $t_{\text{ind}}$ is temperature-dependent because all biosynthetic processes accelerate with temperature, cells at higher temperatures synthesize induction enzymes faster and enter the secondary growth phase sooner. The general form of this temperature dependence can be written as:

$$t_{\text{ind}} = \phi(\mu(\sim, T)) \tag{10}$$

with $\phi$ the decreasing function of $\mu(\sim, T)$. Faster growth at higher temperature shortens the lag.

*Physical interactions.* As cells grow and divide, they occupy space. However, two cells cannot occupy the same location. When a cell grows large enough to divide, the daughter cell must displace its neighbors to find space. This displacement propagates outward through the community, pushing cells away from regions of high local density toward regions of lower density (*31, 32*).

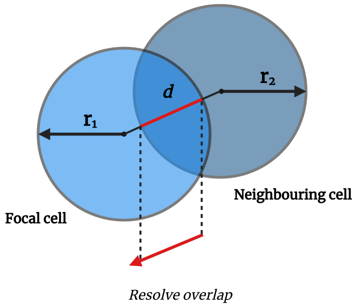


**Figure 6**: **Cell-shoving mechanisms**.

*Endogenous heat production.* When a cell oxidizes a carbon source, only a fraction of the released free energy is captured in ATP and directed toward biosynthesis. The left is dissipated as heat into the immediate surroundings (*7*). Under well-mixed or low-density conditions, this heat dissipates rapidly into the bulk medium and has a negligible effect on cell temperature. In dense, spatially structured communities, however, metabolically active clusters generate heat faster than it can diffuse away, causing local temperature to rise above the ambient bulk value.

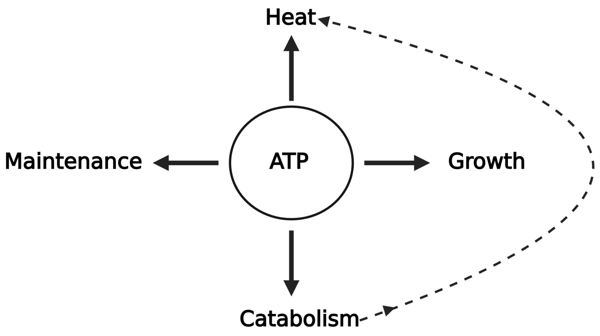


**Figure 7**: **ATP allocation mechanisms**.

*Substrate and metabolite diffusion.* In a spatially structured community, nutrients are not uniformly distributed (*33*). Cells consume substrate locally, depleting it in their immediate vicinity faster than it can be resupplied from the bulk medium. This creates a concentration gradient that substrate is highest far from cells and lowest at the cell surface. The steeper this gradient, the more strongly cells compete for the same pool of nutrients, and the more their growth rate falls below the potential set by bulk concentration alone. Both substrate and metabolite transport are governed by diffusion, in which the net flux of a solute moves from regions of high concentration to regions of low concentration at a rate proportional to the local concentration gradient. The diffusion coefficient $D$ [$\mu m^2$/min] sets the speed of this transport. The time evolution of substrate and metabolite concentrations in the spatial domain is thus described by the reaction-diffusion equations:

$$\frac{\partial S}{\partial t} = D_S \nabla^2 S + R_S(S, \mu(\sim), X) \tag{11}$$

$$\frac{\partial P}{\partial t} = D_P \nabla^2 P + R_P(P, \mu(\sim), X) \tag{12}$$

with $D_S$ and $D_P$ [$\mu m^2$/min] the diffusion coefficients of substrate and metabolite, respectively. $\nabla^2$ the Laplacian operator describing spatial spreading; $R_S$ and $R_P$ the local reaction terms.

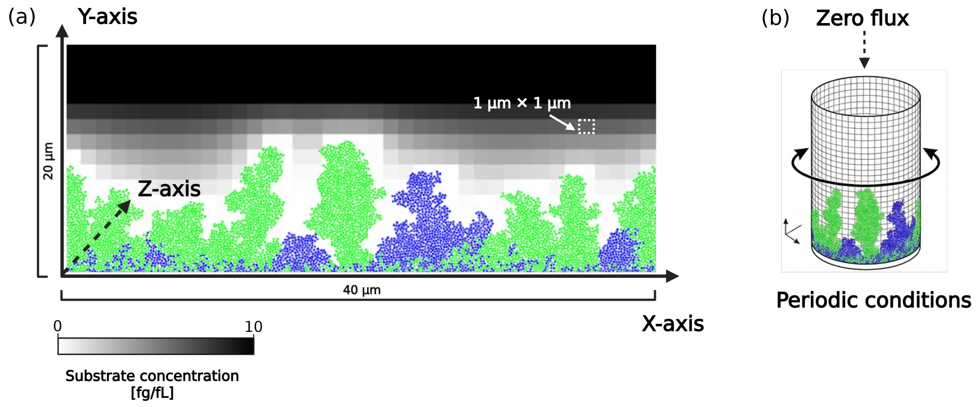


**Figure 8**: **Biofilm simulation environment**.

In the well-mixed culture, any nutrient produced or consumed at one location is immediately redistributed uniformly across the entire community. Concentration gradients cannot form and all cells experience identical substrate, metabolite, and temperature conditions at every moment.

*Heat transfer.* Endogenous heat produced by metabolically active cells enters the local environment as a spatially distributed source. Like nutrients, heat spreads through the medium by diffusion. Thermal energy flows from hot regions toward cooler ones, driven by temperature gradients. The thermal diffusivity $\kappa$[m²/s] sets the speed of nutrient transport. Heat dissipates far more rapidly than molecules diffuse. Thermal diffusivity in aqueous media is approximately $10^3 - 10^4$ times larger than molecular diffusion coefficients (*34*). Local temperature elevation above the ambient bulk value $T_0$ therefore requires either very high local metabolic activity, or sufficiently dense spatial clustering of cells that the collective heat production rate exceeds the rate of thermal dissipation. When such conditions are met, local temperature rises above $T_0$, and the thermal performance curve established earlier begins to amplify metabolism further. The spatial distribution of temperature is governed by the heat equation with a metabolic source term:

$$\frac{\partial T}{\partial t} = \kappa \nabla^2 T + Q_{\text{heat}}(x, y, t) \tag{13}$$

$Q_{\text{heat}}(x, y, t)$ the spatially and temporally varying heat source from local metabolic activity established above.

#### Develop the model

*Temperature-dependent maximum growth rate.* The maximum growth rate of population depends on temperature and is modeled by the Sharpe-Schoolfield equation:

$$r(T) = r_{T_C} \cdot \exp\left[E_a\left(\frac{1}{k \cdot T_C} - \frac{1}{k \cdot T}\right)\right] \cdot \frac{1}{1 + \exp\left[E_h\left(\frac{1}{k \cdot T_h} - \frac{1}{k \cdot T}\right)\right]} \tag{14}$$

with $r(T)$[1/h] the maximum growth rate; $r(T_C)$[1/h] the reference growth rate at reference temperature $T_C$ [K]; $k = 8.62 \times 10^{-5} eV K^{-1}$ the Boltzmann's constant; $E_a$ activation energy; $T$ the temperature in Kelvin (K); $E_h$ the inactivation energy when temperature above the inactivation temperature $T_h$ [K];

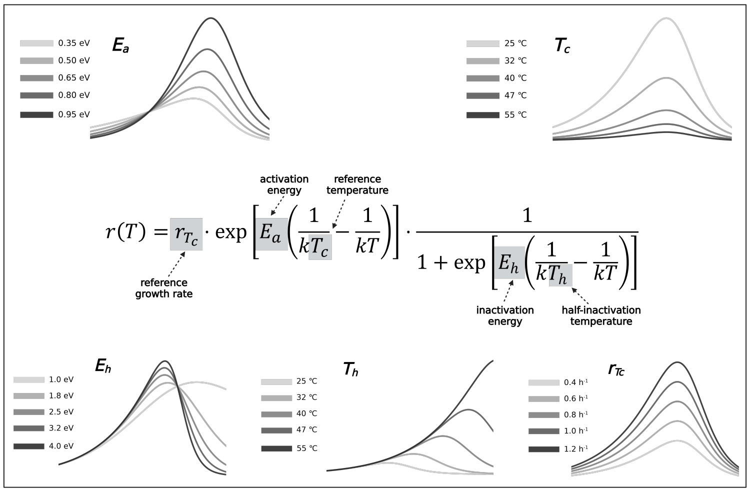


**Figure 9**: **Sharpe-Schoolfield model**.

*Resource-regulated specific growth rate.* The specific growth rate of each cell follows Monod kinetics:

$$\mu_i(\sim) = \begin{cases} r_T \cdot (\frac{S}{K_S+S}) & \text{primary phase} \\ 0 & \text{lag phase} \\ r_T \cdot (\frac{P}{K_P+P}) & \text{secondary phase} \end{cases}$$

with $S$ and $P$ the concentration of primary substrate and secondary metabolite, respectively; $K_S$ and $K_P$ the Monod half-saturation constant for substrate and metabolite, respectively.

*Metabolite production.* The actual carbon allocation is governed by the temperature-dependent bacterial growth efficiency (BGE):

$$\text{BGE}(T) = \frac{Y_{X/n} \cdot \mu}{Y_{X/n} \cdot \mu + m(T)} \tag{15}$$

with $Y_{X/n}$ the pathway-specific biomass yield ($Y_{X/S}$ or $Y_{X/M}$ depending on the active pathway), $\mu$ [1/h] the instantaneous specific growth rate, and $m(T)$ the temperature-dependent maintenance coefficient following Eq. (2). The actual biomass production and $CO_2$ emission per time step are then:

$$\frac{dX}{dt} = \text{BGE}(T) \cdot \frac{\mu}{Y_{X/n}} \cdot X \tag{16}$$

$$\frac{d\text{CO}_2}{dt} = (1 - \text{BGE}(T)) \cdot \frac{\mu}{Y_{X/n}} \cdot X \tag{17}$$

As temperature increases, $m(T)$ rises faster than $\mu(T)$, causing BGE to decline regardless of the active pathway. Consequently, a disproportionately larger fraction of assimilated carbon is respired as $CO_2$ at elevated temperatures. This effect is amplified during metabolite utilization, where the lower biomass yield $Y_{X/M}$ further reduces BGE relative to the substrate pathway, contributing to the steeper temperature dependence of community respiration during the secondary growth phase.

*Diauxic switching.* The transition from primary substrate to secondary metabolite utilization is governed by a three-tier decision process that reflects the biological cost of enzyme synthesis.

Tier 1: Supply-demand evaluation. Cells continuously evaluate whether the local substrate concentration can sustain their growth demands. This is quantified by a supply-demand ratio:

$$\text{Supply-demand ratio} = \frac{\text{Substrate needed}}{\text{Substrate available}} = \frac{S \cdot V_{\text{cell}}}{m_i \cdot (\mu_s / Y_{X/S}) \cdot \Delta t} \tag{18}$$

A supply ratio below a threshold indicates that the local substrate concentration is insufficient to meet maintenance and growth demands, triggering the search for an alternative carbon source. Simultaneously, the local metabolite concentration must exceed a minimum threshold $P > P^*$ to ensure that enzyme synthesis is energetically profitable.

Tier 2: Enzyme induction timer. When both conditions are met, an enzyme induction timer is activated. The induction time is temperature-dependent:

$$t_{\mathrm{ind}}(T) = t_{\mathrm{ind,ref}} \cdot \frac{\mu_{\mathrm{max}}(T_{\mathrm{ref}})}{\mu_{\mathrm{max}}(T)} \tag{19}$$

with $t_{\mathrm{ind,ref}} = 65$ min at $T_{\mathrm{ref}} = 293.15$ K. This representation captures the fact that enzyme synthesis accelerates with metabolic rate at elevated temperatures: warmer cells induce enzymes faster, allowing earlier onset of secondary metabolite utilization.

Tier 3: Pathway selection. Once enzyme induction completes (i.e., $t_{\mathrm{timer}} \geq t_{\mathrm{ind}}$), the cell commits fully to metabolite utilization. The effective growth rate on the secondary metabolite is suppressed by residual primary substrate through catabolite repression:

$$\mu_P^{\mathrm{eff}} = \mu_{\mathrm{max}} \cdot \frac{P}{K_P + P} \cdot \left(1 - \frac{S}{K_{\mathrm{rep}} + S} \cdot \gamma_{\mathrm{rep}}\right) \tag{20}$$

with $K_{\mathrm{rep}}$ the repression half-saturation constant and $\gamma_{\mathrm{rep}}$ the repression strength. This ensures that metabolite utilization is suppressed as long as sufficient primary substrate remains, and only becomes dominant once substrate is effectively depleted. Enzyme status is inherited by daughter cells upon division, providing metabolic memory across generations.

*Physical interactions.* We modeled the physical interactions using disk-shaped particles to calculate vector sums of overlapping radius as below:

$$r_o = s \cdot r_k + r_n - d \tag{21}$$

with $r_o$ [$\mu$m] the sum of overlapping radius, $r_k$ [$\mu$m] and $r_n$ [$\mu$m] the radius length of the focal cell and its neighbor, respectively; d [$\mu$m] the distances between the centres of these two cells, and $s[-]$ the factor to account for a realistic minimum distance between neighboring cells.

*Endogenous heat production.* Heat produced per cell was proportional to its metabolic rate:

$$Q_{\mathrm{heat},i} = \alpha \cdot \mu_i \cdot m_i \tag{22}$$

with $\alpha$ [K.min.fg$^{-1}$] the metabolic heat production coefficient; $\mu_i$ the specic growth rate and $m_i$ the biomass of cell $i$.

**Algorithm 1** Diauxic switching

**Require:** Local substrate $S$, metabolite $P$, cell mass $m_i$, temperature $T$, time step $\Delta t$

$\mu_S \leftarrow \mu_{\max}(T) \cdot S/(K_S + S)$

$S_{\text{needed}} \leftarrow m_i \cdot (\mu_S/Y_{X/S}) \cdot \Delta t$

$\rho \leftarrow S \cdot V_{\text{cell}}/\max(S_{\text{needed}}, \epsilon)$

$t_{\text{ind}}(T) \leftarrow t_{\text{ind,ref}} \cdot \mu_{\max}(T_{\text{ref}})/\mu_{\max}(T)$

**if** $\neg$ `hasEnzyme` **and** $\rho < \rho^*$ **and** $P > P^*$ **then**

  $t_{\text{timer}} \leftarrow t_{\text{timer}} + \Delta t$

  **if** $t_{\text{timer}} \geq t_{\text{ind}}(T)$ **then**

    `hasEnzyme` $\leftarrow$ **true**

  **end if**

**else if** $\rho \geq \rho^*$ **then**

  $t_{\text{timer}} \leftarrow 0$ ▷ Timer resets if substrate recovers

**end if**

*Diffusion process.* Diffusion of substrate and metabolite through the isotropic medium is modeled by the second law of Fick in two dimensions:

$$\frac{\partial S}{\partial t} = D_S \cdot (\frac{\partial^2 S}{\partial x^2} + \frac{\partial^2 S}{\partial y^2}) \tag{23}$$

$$\frac{\partial P}{\partial t} = D_P \cdot (\frac{\partial^2 P}{\partial x^2} + \frac{\partial^2 P}{\partial y^2}) \tag{24}$$

with $S$ and $P$ [fg/fL] the substrate and metabolite concentration, respectively. $D_S$ and $D_P$ [$\mu m^2$/min] the diffusion coefficients of substrate and metabolite, respectively; $x$ and $y$ [$\mu$m] the two spatial dimensions, and $t$ [min] the time. We applied the alternating direction implicit (ADI) scheme to solve the above diffusion PDEs. The ADI implementation consists of two stages:

$$(1 - \frac{D_S}{2} \cdot \frac{\Delta t_1}{\Delta x^2} \cdot \delta_x^2) \cdot S_{(i,j)}(t + \frac{\Delta t_1}{2}) = (1 + \frac{D_S}{2} \cdot \frac{\Delta t_1}{\Delta y^2} \cdot \delta_y^2)) \cdot S_{(i,j)}(t) \tag{25}$$

$$(1 - \frac{D_S}{2} \cdot \frac{\Delta t_1}{\Delta y^2} \cdot \delta_y^2) \cdot S_{(i,j)}(t + \Delta t_1) = (1 + \frac{D_S}{2} \cdot \frac{\Delta t_1}{\Delta y^2} \cdot \delta_x^2)) \cdot S_{(i,j)}(t) \cdot (t + \frac{\Delta t_1}{2}) \tag{26}$$

with $S_{i,j}(t)$ the substrate concentration at time $t$ in the discrete environmental unit with coordinates $(i, j)$, and the second order central differences:

$$\delta_x^2 \cdot S_{(i,j)}(t) = S_{(i+1,j)}(t) - 2S_{(i,j)}(t) - S_{(i-1,j)} \tag{27}$$

$$\delta_y^2 \cdot S_{(i,j)}(t) = S_{(i,j+1)}(t) - 2S_{(i,j)}(t) - S_{(i,j-1)} \tag{28}$$

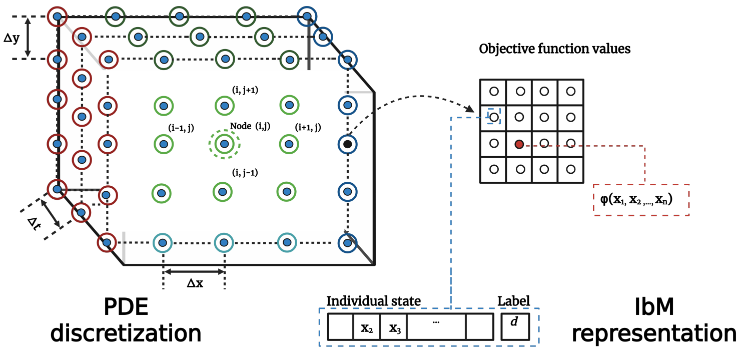


**Figure 10**: **Numerical discretizaiton of diffusion models**.

Heat transfer is modeled by the following heat equation:

$$\frac{\partial T}{\partial t} = \kappa \cdot \left(\frac{\partial^2 T}{\partial x^2} + \frac{\partial^2 T}{\partial y^2}\right) + \sum_{i \in V_{x,y}} \frac{Q_{\text{heat},i}}{\rho \cdot c_\rho \cdot V_{\text{cell}}} \tag{29}$$

with $\kappa$[m$^2$/s] the thermal diffusivity; $\rho \cdot c_\rho$ [J/(m$^3 \cdot K$)] the volumetric heat capacity; $V_{\text{cell}}$ the grid cell volume.

#### Deploy the simulation

*Initialization.* The simulation domain is a $40 \times 20$ $\mu$m$^2$ grid, initialized with a uniform primary substrate concentration of $S_0 = 10.0$ fg/fL and zero metabolite concentrations ($M_A = M_B = 0$ fg/fL). Ten microbial cells (5 Species A, 5 Species B) are placed at random positions within the domain. Each cell is initialized with a biomass drawn from a lognormal distribution, a division factor $\phi = 2 \cdot e^{\mathcal{N}(0,\,0.05)}$ to introduce stochastic variability in division timing, and individual maximum growth rates assigned as:

$$\mu_{\max,i} = \mu_{\max}(T) \cdot (1 + \text{CV} \cdot \mathcal{N}(0, 1)) \tag{30}$$

with coefficient of variation CV $= 0.1$, capturing individual-level variability in growth kinetics. The temperature field is initialized uniformly at the ambient simulation temperature $T_0$, with Dirichlet boundary conditions imposed at the top and bottom boundaries ($T_{\text{boundary}} = T_0$).

*Boundary conditions.* Two sets of boundary conditions are applied depending on the simulation mode. In the spatially structured (biofilm) mode, a zero-flux boundary condition is imposed at the substratum (bottom), and the top boundary is set to a constant substrate concentration ($S_{\text{top}} = 10.0$ fg/fL) to mimic continuous supply from the bulk liquid. Metabolites are set to zero at the top boundary, allowing them to escape the domain. In the well-mixed mode, all spatial gradients are removed by periodically averaging nutrient concentrations across the entire grid every $\Delta t_{\text{mix}} = 0.5$ min, and cell positions are randomly reshuffled at each time step to eliminate spatial structure.

*Scheduling processes.* Biological and physical processes operate on different time scales and are therefore scheduled using a dual time-step scheme. Fast processes — nutrient uptake, metabolic switching, $CO_2$ respiration, and endogenous heat production — are executed at a short time step $\Delta t_1$. Slow processes — cell shoving and spatial overlap resolution — are executed at $\Delta t_2 = 0.1$ min. The slowest process — cell reproduction — is executed at $\Delta t_3 = 0.5$ min. In biofilm mode, $\Delta t_1 = 0.025$ min; in well-mixed mode, $\Delta t_1 = 0.1$ min. The execution order within each time step follows the sequence:

**Algorithm 2** Process schedule per time step

**Require:** Shuffled cell list $C$, time step $\Delta t$

```
for each cell i ∈ C do
    Read local S, M, T from grid
    Update μ_max(T) via Sharpe–Schoolfield (Eq. 2)
    Evaluate enzyme induction (Algorithm 1)
    Select pathway; compute μ_eff, BGE(T)
    Update Δm_i, ΔCO2, Q_heat,i
end for
Diffuse S, M via Fick's second law (Eq. 12)
if ENABLE_TEMPERATURE_FIELD then
    Diffuse T via heat equation (Eq. 14)
end if
if well-mixed mode and t_timer ≥ Δt_mix then
    Homogenize S, M across all grid cells
    Reshuffle all cell positions randomly
end if
```

To avoid artefacts from sequential processing, the cell list is randomly shuffled at each time step before execution. Metabolite decay follows first-order kinetics applied after each diffusion step:

$$M(t + \Delta t) = M(t) \cdot e^{-k_d \Delta t} \tag{31}$$

with decay rate $k_d = 1.667 \times 10^{-5}$ min$^{-1}$. The simulation was implemented in Java using the Repast

Simphony agent-based modelling framework (version 2.x), with spatial neighbor queries accelerated by a KD-tree data structure to efficiently resolve cell shoving interactions.

#### Demonstrate the outputs

*Consumer-resource model.* We adopt the microbial consumer-resource model (MiCRM) to describe the species interactions in a well-mixed culture (*35*). All two symmetrical species share the same growth conditions and consequently will implement diauxic switching simultaneously until all substrates are nearly depleted. The MiCRM consists of four parts: consumer dynamics, resource dynamics, pathway selection, and growth kinetics.

$$\frac{dN_A}{dt} = N_A \cdot \mu_A, \quad \frac{dN_B}{dt} = N_B \cdot \mu_B \tag{32}$$

$$\frac{dS}{dt} = -N_A \cdot I_A^S \cdot \frac{\mu_A^S}{Y_S} - N_B \cdot I_B^S \cdot \frac{\mu_B^S}{Y_S} \tag{33}$$

$$\frac{dP_A}{dt} = N_A \cdot I_A^S \cdot \frac{\alpha}{Y_S} \cdot \mu_A^S - N_B \cdot I_B^P \cdot \frac{1}{Y_P} \cdot \mu_B^{P_A} - k_{\text{decay}} \cdot P_A \tag{34}$$

$$\frac{dP_B}{dt} = N_B \cdot I_B^S \cdot \frac{\alpha}{Y_S} \cdot \mu_B^S - N_A \cdot I_A^P \cdot \frac{1}{Y_P} \cdot \mu_A^{P_B} - k_{\text{decay}} \cdot P_B \tag{35}$$

$$S(0) = S_0, \quad P_A(0) = 0, \quad P_B(0) = 0 \tag{36}$$

$$I_A^S = \mathbf{1}\{\mu_A^S \geq \mu_A^{P_B}\}, \quad I_A^P = 1 - I_A^S \tag{37}$$

$$I_B^S = \mathbf{1}\{\mu_B^S \geq \mu_B^{P_A}\}, \quad I_B^P = 1 - I_B^S \tag{38}$$

$$\mu_A^S = \mu_{\max}^N \cdot r(T) \cdot \frac{S}{K_S + S}, \quad \mu_A^{P_B} = \mu_{\max}^P \cdot r(T) \cdot \frac{P_B}{K_P + P_B} \tag{39}$$

$$\mu_B^S = \mu_{\max}^N \cdot r(T) \cdot \frac{S}{K_S + S}, \quad \mu_B^{P_A} = \mu_{\max}^P \cdot r(T) \cdot \frac{P_A}{K_P + P_A} \tag{40}$$

$$\mu_A = I_A^S \cdot \mu_A^S + I_A^P \cdot \mu_A^{P_B}, \quad \mu_B = I_B^S \cdot \mu_B^S + I_B^P \cdot \mu_B^{P_A} \tag{41}$$

We define the characteristic time for (i) substrate near-depletion as $t^*$ such that $S(t^*) = \varepsilon_S$ with a small tolerance $\varepsilon_S \ll K_S$; and (ii) metabolite near-depletion as $t^{**}$ such that $P_A(t^{**}) + P_B(t^{**}) = \varepsilon_P$ with $\varepsilon_P \ll K_P$. Assume the temperature is uniform for each scenario, so $r(T) = r(T_0)$ is constant.

*Primary phase (from* $0$ *to* $t^*$*).* In primary phase, symmetrical species $A$ and $B$ will only uptake substrate $S$. Thus, we can simplify the Eqs.(32,33,39,40) as below:

$$\frac{dN}{dt} = N \cdot \mu^S(S), \quad \frac{dS}{dt} = -\frac{N \cdot \mu^S(S)}{Y_S}, \quad \mu^S(S) = \mu_{\max}^N \cdot r(T_0) \cdot \frac{S}{K_S + S} \tag{42}$$

We obtain the conservation ratio

$$\frac{dN}{dS} = -Y_S \;\Rightarrow\; N(S) = N_0 + Y_S\,(S_0 - S). \tag{43}$$

Hence

$$\frac{dS}{dt} = -\mu^N_{\max} r(T_0)\,\frac{S}{K_S + S}\left(S_0 + \frac{N_0}{Y_S} - S\right). \tag{44}$$

Integrating from $(0, S_0)$ to $(t, S)$ gives closed formulation for $t^*$

$$t(S) = \frac{1}{\mu^N_{\max} r(T_0)}\,\ln\left[\frac{N_0}{Y_S}\left(\frac{Y_S S_0}{N_0}\right)^{\frac{K_S}{S_0+N_0/Y_S}}\ \frac{\left(S_0 + N_0/Y_S - S\right)^{\frac{K_S}{S_0+N_0/Y_S}-1}}{S^{\frac{K_S}{S_0+N_0/Y_S}}}\right]. \tag{45}$$

Therefore the substrate near-depletion time is

$$t^*(S = \epsilon_S) = \frac{1}{\mu^N_{\max} r(T_0)}\,\ln\left[\frac{N_0}{Y_S}\left(\frac{Y_S S_0}{N_0}\right)^{\frac{K_S}{S_0+N_0/Y_S}}\ \frac{\left(S_0 + N_0/Y_S - \epsilon_S\right)^{\frac{K_S}{S_0+N_0/Y_S}-1}}{S^{\frac{K_S}{S_0+N_0/Y_S}}}\right]. \tag{46}$$

and the population concentration at $t^*$ is

$$N(t^*) = N_0 + Y_S\,(S_0 - \varepsilon_S). \tag{47}$$

During this phase, metabolites are produced but not consumed. Summing Eqs. (34,35) for $P_A, P_B$ yields

$$\frac{d(P_A + P_B)}{dt} = \frac{\alpha}{Y_S}\,(N_A\mu^S_A + N_B\mu^S_B), \qquad \frac{d(P_A + P_B)}{dS} = -\alpha, \tag{48}$$

$$P_A(t^*) + P_B(t^*) = \alpha\,(S_0 - \varepsilon_S). \tag{49}$$

*Metabolite-dominated phase (from $t^*$ to $t^{**}$).* Define the total metabolite $P(t) \equiv P_A(t) + P_B(t)$. At $t^*$,

$$P(t^*) = \alpha\,(S_0 - \varepsilon_S), \qquad N(t^*) = N_0 + Y_S\,(S_0 - \varepsilon_S). \tag{50}$$

Neglecting further production and decay during this phase, we have

$$\frac{dN}{dt} = N\,\mu^P(P), \quad \frac{dP}{dt} = -\frac{N\,\mu^P(P)}{Y_P}, \quad \mu^P(P) = \mu^P_{\max} r(T_0)\,\frac{P}{K_P + P}. \tag{51}$$

Thus

$$\frac{dN}{dP} = +Y_P \;\Rightarrow\; N(P) = N(t^*) + Y_P\big(P(t^*) - P\big). \tag{52}$$

Consequently

$$\frac{dP}{dt} = -\mu^P_{\max} r(T_0)\,\frac{P}{K_P + P}\left(P(t^*) + \frac{N(t^*)}{Y_P} - P\right). \tag{53}$$

Integrating from $(t^*, P(t^*))$ to $(t, P)$ yields

$$t^{**} = t^* + \frac{1}{\mu^P_{\max} r(T_0)}\,\ln\left[\frac{N(t^*)}{Y_P}\left(\frac{Y_P\,P(t^*)}{N(t^*)}\right)^{\frac{K_P}{P(t^*)+N(t^*)/Y_P}}\ \frac{\big(P(t^*) + N(t^*)/Y_P - P\big)^{\frac{K_P}{P(t^*)+N(t^*)/Y_P}-1}}{P^{\frac{K_P}{P(t^*)+N(t^*)/Y_P}}}\right]. \tag{54}$$

*Total $CO_2$ (Primary + Secondary).* Let $C_S, C_P$ be carbon per unit $S, P$ and $Y^C_S, Y^C_P$ the carbon yields to biomass (biomass C formed per resource C consumed). Let $\kappa_S, \kappa_P$ be carbon-to-$CO_2$ factors (typically

$\kappa_S \simeq \kappa_P \simeq 1$), and define the substrate-to-metabolite carbon conversion $\alpha_C \equiv \alpha\,(C_P/C_S)$. Then the substrate-phase and metabolite-phase cumulative $CO_2$ are

$$CO_2{}^{(S)} = \kappa_S\,[1 - Y_S^C - \alpha_C]\; C_S\,(S_0 - \varepsilon_S), \qquad CO_2{}^{(P)} = \kappa_P\,[1 - Y_P^C]\; C_P\,\big(P_A(t^*) + P_B(t^*) - \varepsilon_P\big). \tag{55}$$

Using $P_A(t^*) + P_B(t^*) = \alpha\,(S_0 - \varepsilon_S)$, the total is

$$CO_2{}^{(\text{total})} = \kappa_S[1 - Y_S^C - \alpha_C]\,C_S\,(S_0 - \varepsilon_S) + \kappa_P[1 - Y_P^C]\,C_P\,\big(\alpha\,(S_0 - \varepsilon_S) - \varepsilon_P\big). \tag{56}$$

If $S, P$ are measured in equivalent carbon units ($C_S = C_P = 1$), then $\alpha_C = \alpha$ and

$$CO_2{}^{(\text{total})} = [1 - Y_S^C - \alpha]\,(S_0 - \varepsilon_S) + [1 - Y_P^C]\,\big(\alpha\,(S_0 - \varepsilon_S) - \varepsilon_P\big). \tag{57}$$

From the above-derived formulations, we can further analyze how the temperature response of microbial respiration will be affected.

#### Characteristic time scale as $[r(T)]^{-1}$

Both characteristic times $t^*$ and $t^{**}$ derived in the previous section take the form

$$t^*(T) = \frac{1}{\mu_{\max}^N \cdot r(T)} \cdot F(N_0, S_0, Y_S, K_S), \qquad t^{**}(T) - t^*(T) = \frac{1}{\mu_{\max}^P \cdot r(T)} \cdot G(N^*, P^*, Y_P, K_P) \tag{58}$$

with functions $F$ and $G$ separate the stoichiometric and half-saturation constants from temperature factors. Both characteristic times decrease with warming at the same rate $r(T)$, and their ratio $t_{\text{ind}}(T)/[t^{**}(T) - t^*(T)]$ is temperature-independent:

$$\frac{t_{\text{ind}}(T)}{t^{**}(T) - t^*(T)} = \frac{t_{\text{ind,ref}} \cdot \mu_{\max}(T_{\text{ref}})}{\mu_{\max}(T)} \cdot \frac{\mu_{\max}(T)}{G} = \frac{t_{\text{ind,ref}} \cdot \mu_{\max}(T_{\text{ref}})}{G} = \text{const.} \tag{59}$$

This formulation indicates that whether the induction lag fits within the secondary phase window is determined by stoichiometric parameters alone, not by temperature. The temperature-dependent amplification of $E_a$ therefore does not arise from a simple threshold crossing in time. Instead, the below coupled mechanisms drive the amplification.

#### BGE links maintenance to apparent $E_a$

From Eqs.(16,17), the instantaneous community respiration rate during either phase is

$$R_n(T) = \big[1 - \text{BGE}_n(T)\big] \cdot \frac{\mu(T)}{Y_{X/n}} \cdot X, \qquad n \in \{S, P\} \tag{60}$$

with $\mathrm{BGE}_n(T) = Y_{X/n}\,\mu(T)/[Y_{X/n}\,\mu(T) + m(T)]$. In the Boltzmann limit (ignoring the high-$T$ Schoolfield correction), $\mu(T) \propto e^{-E_a/kT}$ and $m(T) \propto e^{-E_m/kT}$. Substituting into Eq. (60) gives

$$\begin{aligned} R_n(T) &\propto \frac{\mu(T)\cdot m(T)}{Y_{X/n}\,\mu(T) + m(T)} \\ &\propto \frac{e^{-(E_a+E_m)/kT}}{Y_{X/n}\,e^{-E_a/kT} + e^{-E_m/kT}} \end{aligned} \tag{61}$$

Taking the logarithmic derivative with respect to $1/kT$ yields the *apparent activation energy*:

$$E_a^{\mathrm{app},n} = E_a \cdot \mathrm{BGE}_n(T) + E_m \cdot [1 - \mathrm{BGE}_n(T)] \tag{62}$$

This is a weighted average of $E_a$ (the kinetic activation energy) and $E_m$ (the maintenance activation energy), with weight $1 - \mathrm{BGE}_n$ increasing with temperature as $m(T)$ grows faster than $\mu(T)$ (since $E_m > E_a$). Equation (62) has two implications. First, $E_a^{\mathrm{app}}$ is bounded between $E_a$ and $E_m$ at all temperatures, as it approaches $E_a$ when growth dominates maintenance ($Y_{X/n}\,\mu \gg m$) and approaches $E_m$ when maintenance dominates. Second, because $Y_{X/P} < Y_{X/S}$, the metabolite pathway operates at a lower BGE than the substrate pathway at every temperature, so $E_a^{\mathrm{app},P} > E_a^{\mathrm{app},S}$ holds universally.

#### Community $E_a$ amplified by mutualism.

The total community respiration integrates both phases:

$$R_{\mathrm{total}}(T) = \phi_S(T)\cdot R_S(T) + \phi_P(T)\cdot R_P(T) \tag{63}$$

where $\phi_S(T)$ and $\phi_P(T)$ are the time-weighted fractions of substrate and metabolite utilization. In the *baseline* scenario (no mutualism), $\phi_P = 0$ and $E_a^{\mathrm{community}} = E_a^{\mathrm{app},S}$. When mutualism is enabled, the onset of the secondary phase depends on whether the induction lag $t_{\mathrm{ind}}(T)$ is short relative to the *batch observation window* $t_{\mathrm{obs}}$:

$$\phi_P(T) = \max\left(0,\ \frac{t_{\mathrm{obs}} - t^*(T) - t_{\mathrm{ind}}(T)}{t_{\mathrm{obs}}}\right) \tag{64}$$

Since $t^*(T) \propto [r(T)]^{-1}$ decreases with $T$ while $t_{\mathrm{obs}}$ is fixed, $\phi_P(T)$ is an increasing function of temperature once $t^*(T) + t_{\mathrm{ind}}(T) < t_{\mathrm{obs}}$. Combined with corollary (2) above ($E_a^{\mathrm{app},P} > E_a^{\mathrm{app},S}$), the community apparent activation energy

$$E_a^{\mathrm{community}}(T) = \frac{d\ln R_{\mathrm{total}}}{d(1/kT)} = \phi_S(T)\cdot E_a^{\mathrm{app},S} + \phi_P(T)\cdot E_a^{\mathrm{app},P} + \frac{d\phi_P}{d(1/kT)}\cdot \ln\frac{R_P}{R_S} \tag{65}$$

exceeds $E_a^{\mathrm{app},S}$ whenever (i) $\phi_P(T) > 0$ and (ii) $E_a^{\mathrm{app},P} > E_a^{\mathrm{app},S}$, both satisfied here. The third term (the phase-fraction gradient) provides additional super-Arrhenius amplification in the temperature range where $\phi_P(T)$ is rising steeply. This is the mechanistic origin of the $E_a$ increase from 0.65 eV (baseline) to 0.77 eV (mutualism) reported in Fig. 2.

#### Spatial structure and endogenous heat as further amplifiers

The analytical framework above is derived for a well-mixed, spatially homogeneous system. In the IbM, two additional mechanisms further amplify the apparent $E_a$. In the biofilm, the metabolite concentration available to cell $i$ is not the community average but the *local* value governed by diffusion. The local penetration depth of metabolite scales as (see Section S4):

$$\ell(T) \propto \left[\frac{D_P}{\mu_{\max}(T)}\right]^{1/2} \propto [r(T)]^{-1/2} \tag{66}$$

Warming reduces $\ell(T)$, steepening metabolite gradients and concentrating cross-feeding activity into smaller but more metabolically intense hotspots. Simultaneously, faster cluster expansion at higher $T$ reduces the physical separation between partner species. Both effects cause $\phi_P(T)$ to rise more steeply than in Eq. (64), producing the additional 53% amplification (i.e., $E_a : 0.77 \to 1.18$ eV).

When endogenous heat production is enabled, local temperature $T_{\text{local}} = T_0 + \Delta T_i$ feeds back on $\mu_{\max}(T_{\text{local}})$ and hence on $R_P$. The local temperature excess scales as:

$$\Delta T_i \propto \frac{\alpha \cdot \mu_i \cdot m_i}{\kappa/\ell_{\text{heat}}^2} \propto \mu_{\max}(T) \tag{67}$$

Because $\Delta T_i$ grows with ambient $T$, the effective local $r(T_{\text{local}})$ exceeds $r(T_0)$ by a margin that itself increases with $T_0$. This positive feedback ($T \to \mu \to Q_{\text{heat}} \to T$) is the origin of the additional amplification (i.e., $E_a : 1.18 \to 1.78$ eV). Together, the three mechanisms amplify the apparent activation energy in a sequential and mechanistically grounded manner:

$$E_a^{\text{community}} = \underbrace{E_a + (E_m - E_a)(1 - \overline{\text{BGE}})}_{\text{BGE mechanism}} + \underbrace{\delta_\phi}_{\text{mutualistic phase shift}} + \underbrace{\delta_\ell}_{\text{spatial threshold}} + \underbrace{\delta_{\Delta T}}_{\text{heat feedback}} \tag{68}$$

where each $\delta > 0$ and is zero in the absence of the corresponding mechanism.

#### Validation across scenarios

We validate the amplification mechanisms across different thermal performance scenarios.

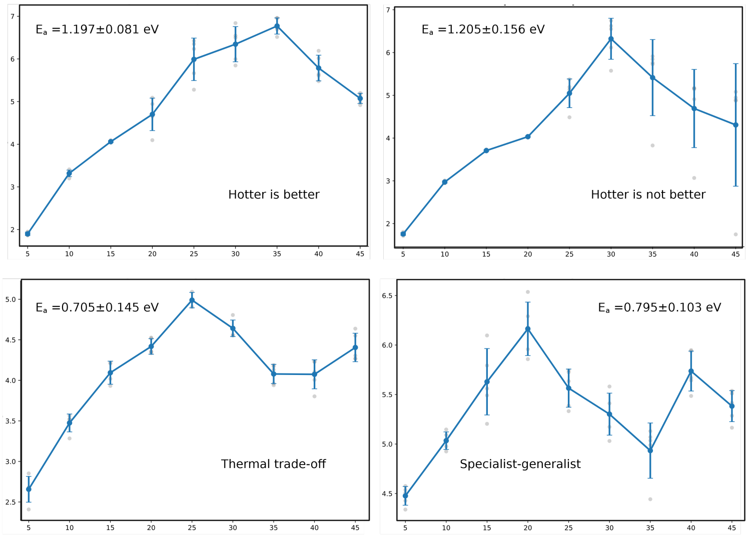


**Figure 11**: **The temperature sensitivity of microbial respiration in biofilm culture with different TPC scenarios**

and Monod kinetics scenarios

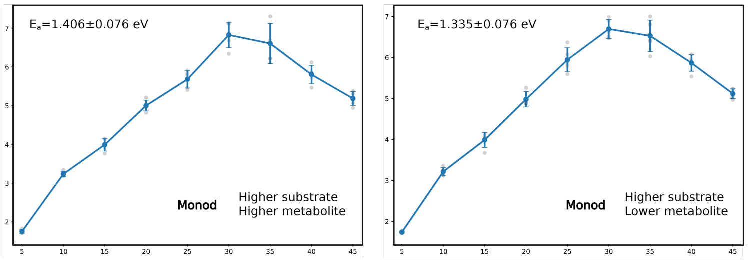


**Figure 12**: **The temperature sensitivity of microbial respiration in biofilm culture with different Monod kinetics scenarios**

**Table 1**: **Thermal performance curve parameters.** Both species share identical Sharpe–Schoolfield parameters in the symmetrical scenario used throughout the main text. $k = 8.62 \times 10^{-5}$ eV $K^{-1}$ is the Boltzmann constant.

| Symbol | Description | Value [unit] |
|---|---|---|
| $E_a$ | Activation energy (Arrhenius term) | 0.65 [eV] |
| $E_h$ | Thermal inactivation energy | 2.5 [eV] |
| $T_h$ | Half-inactivation temperature | 310.15 [K] (37 °C) |
| $T_C$ | Reference temperature for $r_{T_C}$ | 293.15 [K] (20 °C) |
| $r_{T_C}$ | Reference maximum growth rate at $T_C$ | 0.8 [$h^{-1}$] |
| $E_m$ | Activation energy of maintenance metabolism | 0.75 [eV] |
| $E_{h,m}$ | High-temperature inactivation energy of maintenance | 3.0 [eV] |
| $T_m$ | Half-inactivation temperature for maintenance | 323.15 [K] (50 °C) |
| $T_0$ | Simulated ambient temperature range | 5–45 [°C], step 5 °C |

**Table 2**: **Resource kinetics and carbon stoichiometry parameters.** Monod half-saturation constants and yield coefficients for primary-substrate and secondary-metabolite pathways. Concentrations are expressed in fg fL$^{-1}$ (equivalent to $\mu$g mL$^{-1}$).

| Symbol | Description | Value [unit] |
|---|---|---|
| *Monod kinetics* | | |
| $K_S$ | Half-saturation constant for primary substrate (Species A & B) | $4.3 \times 10^{-4}$ [fg fL$^{-1}$] |
| $K_{P,A}$ | Half-saturation constant for metabolite A (consumed by Species A) | $3.6 \times 10^{-4}$ [fg fL$^{-1}$] |
| $S_0$ | Initial primary substrate concentration | 10.0 [fg fL$^{-1}$] |
| *Carbon stoichiometry — primary substrate pathway* | | |
| $Y_{X/S}$ | Biomass yield per unit substrate consumed | 0.51 [fg fg$^{-1}$] |
| $Y_{P/S}$ | Metabolite yield per unit substrate consumed (overflow) | 0.19 [fg fg$^{-1}$] |
| $Y_{CO_2/S}$ | $CO_2$ yield per unit substrate consumed | 0.49 [fg fg$^{-1}$] |
| *Carbon stoichiometry — secondary metabolite pathway* | | |
| $Y_{X/M}$ | Biomass yield per unit metabolite consumed | 0.29 [fg fg$^{-1}$] |
| $Y_{CO_2/M}$ | $CO_2$ yield per unit metabolite consumed | 1.03 [fg fg$^{-1}$] |
| *Maintenance respiration* | | |
| $m_{\text{ref}}$ | Baseline maintenance coefficient at $T_C$ | 0.020 [h$^{-1}$] |
| *Overflow and decay* | | |
| $\alpha$ | Base overflow fraction (substrate $\rightarrow$ metabolite at $T_{\text{ref}}$) | 0.50 [fg fg$^{-1}$] |
| $Q_{10}^{\text{ov}}$ | Temperature coefficient for overflow metabolism | 2.0 [–] |
| $k_d$ | First-order metabolite decay rate | $1.667 \times 10^{-5}$ [min$^{-1}$] |

**Table 3**: **Diauxic switching and enzyme induction parameters.** These govern the three-tier substrate-to-metabolite transition described in Algorithm 1.

| Symbol | Description | Value [unit] |
|---|---|---|
| $t_{\text{ind,ref}}$ | Enzyme induction lag at reference temperature $T_{\text{ref}}$ | 265 [min] |
| $T_{\text{ref}}$ | Reference temperature for induction scaling | 278.15 [K] (5 °C) |
| $\rho^*$ | Supply–demand ratio threshold triggering induction | 0.005 [–] |
| $P^*$ | Minimum metabolite concentration for induction to begin | 0.003 [fg fL$^{-1}$] |
| $K_{\text{rep}}$ | Half-saturation constant for catabolite repression | 0.001 [fg fL$^{-1}$] |
| $\gamma_{\text{rep}}$ | Catabolite repression strength | 0.95 [–] |
| $\mu_{\max}^{P}$ | Metabolite-pathway maximum growth rate scaler | $1.0 \times \mu_{\max}(T)$ [–] |

**Table 4**: **Simulation domain, cell properties, and numerical parameters.** All spatial quantities are in micrometres ($\mu$m) and temporal quantities in minutes (min) unless stated otherwise.

| Symbol | Description | Value [unit] |
|---|---|---|
| *Spatial domain* | | |
| $L_x \times L_y$ | Simulation grid dimensions | $40 \times 20$ [$\mu$m$^2$] |
| $\Delta x$ | Grid cell width | 10 [$\mu$m] |
| $\Delta y$ | Grid cell height | 10 [$\mu$m] |
| $V_{\text{cell}}$ | Grid cell volume | 1000 [$\mu$m$^3$] |
| *Diffusion and heat transfer* | | |
| $D_S$ | Diffusivity of primary substrate | 0.1 [$\mu$m$^2$ min$^{-1}$] |
| $D_P$ | Diffusivity of secondary metabolite | 10.0 [$\mu$m$^2$ min$^{-1}$] |
| $\kappa$ | Thermal diffusivity | 200.0 [$\mu$m$^2$ min$^{-1}$] |
| $k_{\text{loss}}$ | Heat loss rate to surroundings | $1.0 \times 10^{-5}$ [min$^{-1}$] |
| $\alpha_Q$ | Metabolic heat production coefficient | 300.0 [K min fg$^{-1}$] |
| *Temporal discretisation* | | |
| $\Delta t_1$ | Fast time step (nutrient uptake, diffusion) | 0.025 [min] (biofilm); 0.1 [min] (well-mixed) |
| $\Delta t_2$ | Slow time step (cell shoving) | 0.1 [min] |
| $\Delta t_3$ | Reproduction time step | 0.5 [min] |
| $\Delta t_{\text{mix}}$ | Well-mixed homogenisation period | 0.5 [min] |
| *Cell properties* | | |
| $\rho_{\text{cell}}$ | Cell density | 290.0 [fg $\mu$m$^{-3}$] |
| $s$ | Shoving parameter (minimum distance factor) | 1.3 [–] |
| CV | Coefficient of variation for individual $\mu_{\max}$ | 0.1 [–] |
| $N_0$ | Initial number of cells (each species) | 5 [cells] |

22. S. Arrhenius, On the heat of dissociation and the influence of temperature on the degree of dissociation of the electrolytes. *Zeitschrift für Physikalische Chemie (in German, Über die Dissociationswärme und den Einfluss der Temperatur auf den Dissociationsgrad der Elektrolyte)* **4** (1), 96–116 (1889).
